# Post-inhibitory rebound firing drives hypothalamic activation for male mating

**DOI:** 10.1101/2025.08.25.672248

**Authors:** Wen Zhang, Xi Zha, Liping Gong, Liu Fan, Zhitao Feng, Zhewei Zhang, Xuanzi Cao, Yunxian Bai, Zhuolei Jiao, Shuaishuai Li, Ziyue Wang, Xiaojing Ding, Tianming Yang, Huatai Xu, Yuanyuan Mi, Xiaohong Xu

**Author notes:** Correspondence (Y.M.); (X.X.). These authors contributed equally (Wen Zhang, Xi Zha). These authors contributed equally (Liping Gong, Liu Fan, Zhitao Feng, Zhewei Zhang, Xuanzi Cao, Yunxian Bai).

## Abstract

Many behaviorally relevant limbic circuits are dominated by inhibitory connections, raising the question of how neuronal activation arises in such contexts. In male mating, both activation of mPOA and its inhibitory inputs are essential—a paradox previously ascribed to disinhibition. Here, we show that the mPOA largely lacks local inhibitory circuits, arguing against disinhibition. Instead, *in vivo* recordings reveal that mount-active mPOA neurons are suppressed prior to behavioral onset, and this inhibition negatively correlates with subsequent activation, consistent with post-inhibitory rebound firing (PIR). A biologically plausible conductance-based model demonstrates that synaptic inhibition interacts with T-type Ca²⁺ currents — the ionic basis of PIR — to promote mPOA neuron firing. Experimentally, bidirectional manipulations of mPOA T-type Ca²⁺ channels confirm this mechanism: knockdown reduces PIR and impairs male mating, while overexpression enhances mating. Together, these results identify PIR as a critical mechanism by which inhibition promotes activation in limbic circuits to drive male mating.

## Introduction

Male mice exhibit a stereotyped mating sequence that progresses from appetitive social and chemoinvestigation to consummatory mounting and rhythmic pelvic thrusting^1,2^. This behavioral progression is orchestrated by the medial preoptic area (mPOA), an evolutionarily conserved brain region^3,4^. In mice, mPOA neuronal activity ramps before mounting and thrusting^3,5,6^. Artificial activation of the mPOA triggers mounting and thrusting, even toward inappropriate targets, whereas its inhibition disrupts consummatory mating^3,5^. Notably, under normal mating conditions, the transition from social investigation to mounting is variable and unpredictable despite the constant presence of a receptive female, suggesting that mPOA neural dynamics, rather than sensory cues alone, govern the initiation of consummatory mating.

The mPOA integrates both excitatory and inhibitory inputs from upstream limbic structures^2^. Among these, excitatory input from the posterior amygdala (PA) is active throughout mating and is essential for mating progression^7^. Chemogenetic inhibition of PA neurons that project to mPOA prolongs appetitive chemoinvestigation and nearly abolishes mounting and pelvic thrusts, indicating excitatory PA input is necessary for the transition to consummatory behavior. However, optogenetic stimulation of PA neurons alone fails to elicit mounting^7^, suggesting that while PA input is necessary, it is insufficient to drive the mPOA activity patterns required for mounting initiation — unlike direct optogenetic activation of the mPOA^3^.

In parallel, the mPOA receives strong inhibitory input from the principal nucleus of the bed nucleus of the stria terminalis (BNSTpr)^5,8^, which is also active during mating^5,6,9^. Chemogenetic inhibition of BNSTpr neurons or acute silencing of BNSTpr → mPOA projections during female approach abolishes the transition to mounting^6,10^, underscoring a critical role for inhibitory inputs in shaping mPOA activity at mounting onset. Moreover, optogenetic activation of BNSTpr neurons promotes inappropriate male-male mounting, indicating their sufficiency to bypass sensory constraints to drive mating^10^.

The BNSTpr-derived neuropeptide, Tac1, facilitates long-term potentiation (LTP) of excitatory synapses onto mPOA neurons^5^, enabling a gradual buildup of excitability over minutes. However, this form of plasticity could not fully account for the rapid, second-scale activity changes observed at the onset of mounting — particularly given that, as noted above, excitatory inputs alone were insufficient^7^, whereas inhibitory synaptic inputs were required^3,6^, to generate the precise temporal dynamics underlying mounting initiation in the mPOA. To explain this, a disinhibition model has been proposed^6,11^, in which BNST-derived inhibition is thought to suppress local inhibitory interneurons, thereby activating mPOA mount-related neurons and promoting mating progression. Yet, direct evidence for a local inhibitory microcircuit within the mPOA remains lacking.

Here, we performed *ex vivo* and *in vivo* recordings and found little evidence of local connectivity among mPOA neurons, arguing against disinhibition as the mechanism for inhibition-driven activation. Instead, *in vivo* single-unit recordings revealed that mounting-active mPOA neurons receive strong inhibition prior to behavior onset, which negatively correlated with subsequent neuronal activation during mounting. This observation, together with previous *ex vivo* evidence that a subset of mPOA neurons exhibits post-inhibitory rebound firing (PIR)^12^, led us to investigate the contribution of PIR to mPOA neural dynamics underlying male mating. Using mathematical modeling, gene manipulation, and behavioral analysis, we suggest that synaptic inhibition synergizes with T-type Ca²⁺ currents to promote mPOA neuron firing, establishing PIR as a critical mechanism linking inhibition to mPOA activation to drive mating progression in males.

## Results

### Lack of local inhibitory microcircuits in the mPOA

To determine whether the mPOA contains local microcircuits, we performed paired whole-cell recordings from neighboring neurons in acute coronal or sagittal brain slices (**Figure 1A-1C**), following established methods^13–15^. Briefly, action potentials were evoked in one neuron while a nearby neuron was voltage-clamped at – 40 mV to detect potential monosynaptic connections (**Figure 1B**). As the positive control, we applied this protocol to the barrel cortex (Figure S1A-S1B) and tested 20 pairs of connections: 12 excitatory-to-excitatory (E→E), 4 excitatory-to-inhibitory (E→I), and 4 inhibitory-to-inhibitory (I→I), with putative cell types identified by soma morphology. Out of these tested pairs in the barrel cortex, 3 of the 4 I→I connections exhibited synaptic responses, confirming sensitivity of the method (Figure S1C-S1D). In contrast, none of the 137 tested pairs in the mPOA showed synaptic responses (**Figure 1D**), consistent with previous reports of sparse local connectivity in hypothalamic regions^15^.

**Figure 1.**
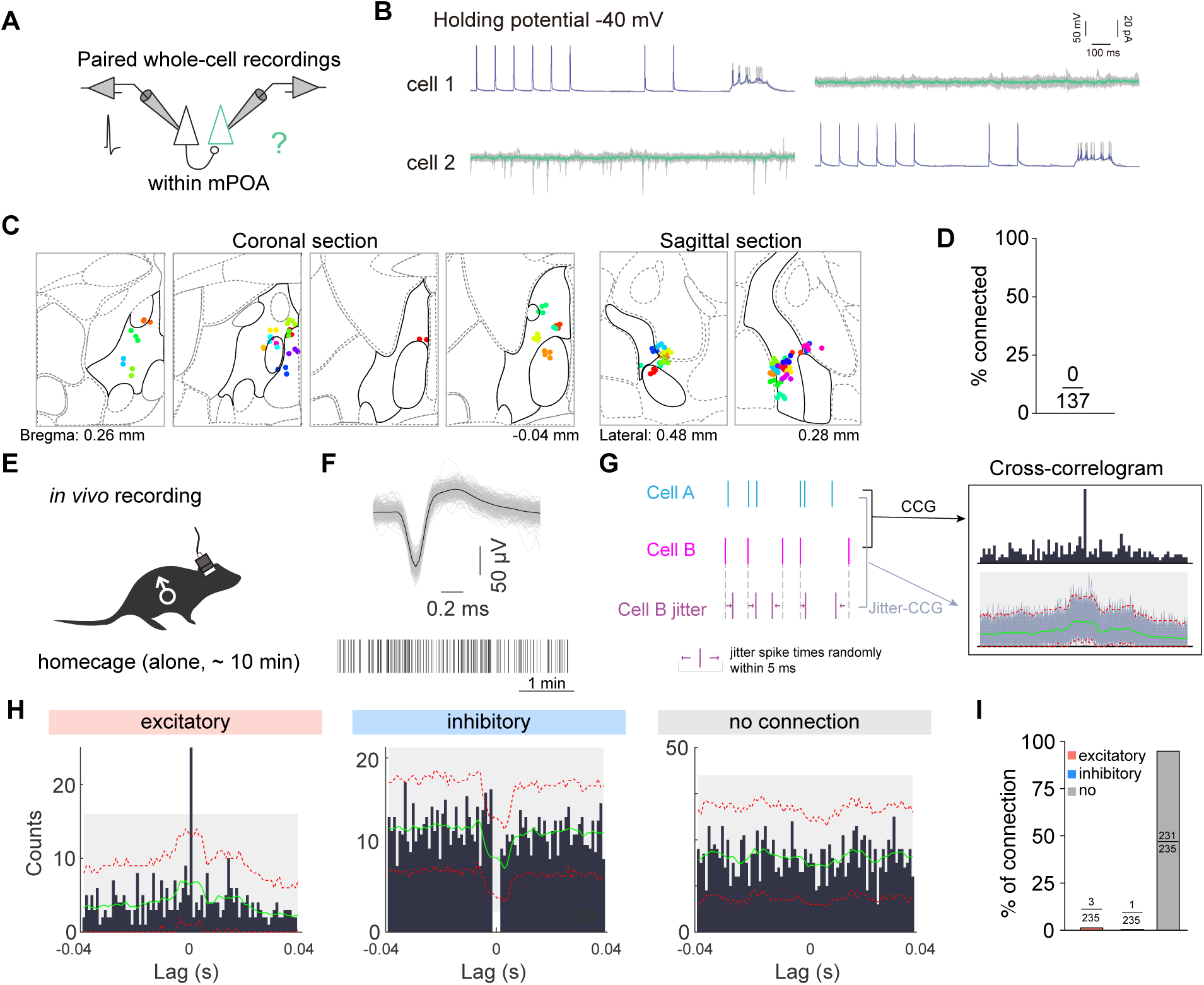
*Ex vivo* and *in vivo* recordings reveal the near absence of local inhibitory microcircuits in the mPOA. **(A–D) *Ex vivo* recordings in the mPOA.** (A). Schematic of paired whole-cell recordings from mPOA neurons in acute brain slices prepared in both coronal and sagittal planes. (B). Representative traces from paired *ex vivo* recordings, showing no synaptic connectivity between mPOA neurons. Blue traces indicate the averaged membrane potential in the stimulated cell with evoked action potentials; green traces indicate the averaged post-synaptic responses in the unstimulated cell, held at -40 mV. The original raw traces are shown in gray. (C). Anatomical locations of all recorded neurons mapped onto reference coronal and sagittal brain atlas sections from the Allen Brain Atlas (https://mouse.brain-map.org/static/atlas), with the mPOA outlined in each section by thick black lines. (D). Quantification of synaptically connected neuron pairs. No evidence of monosynaptic connectivity was detected among the 137 tested mPOA → mPOA pairs. N = 53 pairs from 6 males in the coronal plane and 84 pairs from 3 males in the sagittal plane. **(E- F) *In vivo* single-unit recordings from the mPOA of freely moving male mice in their homecage.** (E) Schematic of recording setup. (F) Representative spike waveform (top) and raster plot (bottom) from an isolated unit. **(G-I) Jitter-based cross-correlogram (CCG) analysis.** (G) The true correlogram (top) of two cells’ spike timing is compared to correlograms generated from jittered spike trains that preserve overall firing rates but randomize the precise timing, enabling detection of significant short-latency interactions. (H) Example CCGs illustrating three types of functional relationships: excitatory (left), inhibitory (middle), and no connection (right). Each histogram shows the spike pair counts (black bars) as a function of time lag (± 40 ms) between a reference neuron and a target neuron over 5 minutes of recording. The green line denotes the mean of the jittered CCGs distribution; red lines mark the 0.5% and 99.5% confidence levels; and gray shading indicates the global confidence band. A short-latency (< 4 ms) peak or trough exceeding the global confidence bands indicates a putative monosynaptic excitatory or inhibitory connection, respectively. (I). Summary of the proportion of excitatory, inhibitory, and unconnected pairs within the mPOA. N = 235 pairs from 6 males. *See also Figures S1 and S2A-2B*.

Because *ex vivo* paired recordings are spatially limited to adjacent neurons, we next assessed functional local connectivity *in vivo*. We implanted electrodes into the mPOA and recorded single-unit activity from freely behaving, sexually experienced male mice alone in their home cages, in the absence of any social stimuli (**Figure 1E- 1F**, Figure S2A-S2B). To detect potential functional local interactions among mPOA neurons, we analyzed baseline activity (alone in homecage for 5 min) of simultaneously recorded units for short-latency correlation or anticorrelations (**Figure 1G**). Across 235 valid single-unit pairs recorded from six C57BL/6 males, only one pair showed short-latency anticorrelation suggestive of local inhibition, and three pairs showed short-latency correlation indicative of local excitation (**Figure 1H-1I**). The remaining pairs showed no significant interaction (**Figure 1I**). Thus, these *in vivo* results, consistent with the *ex vivo* findings, demonstrating that the mPOA lacks robust local microcircuits, challenge the proposed disinhibitory mechanism of mPOA activation during male mating^6,7,11^.

### Single-unit recordings reveal inhibitory gating of mPOA mount-activated neurons

To further examine mPOA neural dynamics underlying mating, particularly around the onset of mounting, we recorded single-unit activity in male mice interacting with hormonally primed, receptive females **(Figure 2A**). As described previously^3^, male mating behavior progresses through distinct phases: beginning with appetitive behaviors such as social investigation (SI) and anogenital sniffing (sniff), followed by consummatory behaviors including mounting and pelvic thrusting that culminate in ejaculation **(Figure 2B**). To identify behaviorally relevant neurons, we normalized each unit’s firing rate to baseline periods devoid of social interactions or mating attempts and compared activity during each annotated behavior to the corresponding 5 – 10 s pre-onset window (**Figure 2B**; see Methods). Units showing significant activity changes (p < 0.05, paired t-test) were classified as activated or inhibited based on the direction of change (Figure S2C–S2G). Notably, units responsive to mounting and thrust exhibited stronger behavioral modulation than those responsive to SI or sniffing (Figure S2C– S2G), consistent with prior population-level findings^3^. Moreover, there was substantial overlap between mount- and thrust-responsive units, whereas SI- and sniff-responsive units showed no significant overlap with each other (Figure S2H), in agreement with previous Ca²⁺ imaging studies^6^.

**Figure 2.**
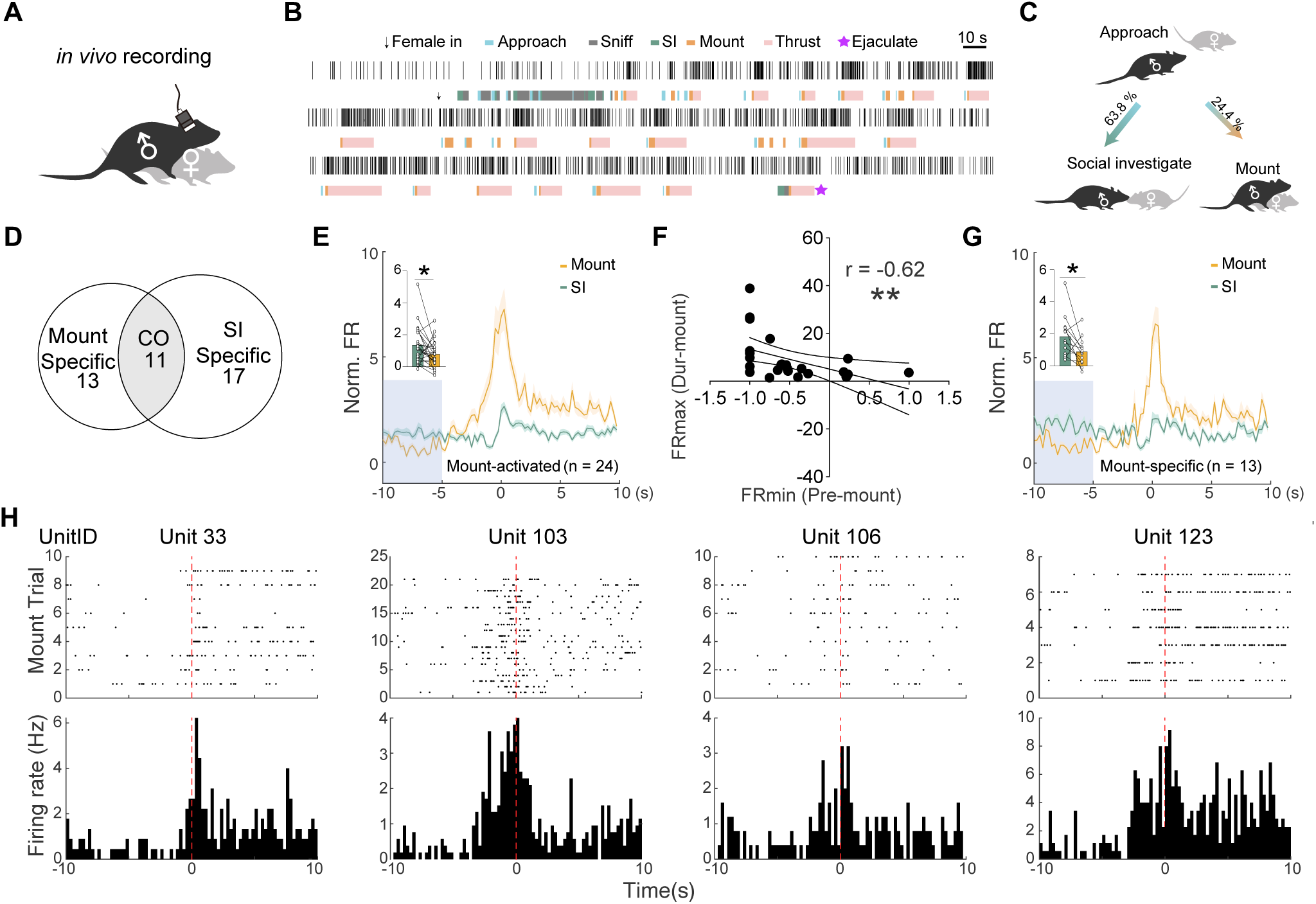
Inhibitory gating of mount-activated units in the mPOA (A) Schematic of *in vivo* single-unit recordings from the mPOA in freely moving male mice mating with a receptive female in their homecage. N = 6 males. (B) Example raster plots of a single-unit with behavioral annotations shown below. (C) Illustration of the transition from approach to social investigation (SI) or mounting. (D) Venn diagram showing the number of units selectively activated during mounting (Mount specific), social investigation (SI specific), or both (CO). (E) Normalized firing rates (FR) of all mount-activated units (N = 24) aligned to the onset (time 0) of mount (orange) or SI (green) bouts. Shaded areas indicate SEM. A reduction in firing preceding mount bouts (gray box), relative to SI bouts, is quantified in the inset bar plot. (F) Correlation between the trough normalized FR (FRmin) in the pre-mount period (gray box in E) and the peak normalized FR (FRmax) during mounting for all mount- activated units (N=24). A significant negative correlation is observed. (G) Normalized FR of all mount-specific neurons (N=13) aligned to the onset of mount (orange) or SI (green) bouts. Pre-mount inhibition (gray box) is quantified in the inset bar plot. (H) Raster plots (top) and peri-event time histogram (PETH; bottom) from four representative mount-activated neurons, showing a PIR-like firing pattern characterized by prolonged suppression followed by rapid excitation. In the raster plots, each row represents a single mount event aligned to onset at time 0. *p < 0.05, **p < 0.01. *See also Figures S2*.

Focusing on the transition from the appetitive to the consummatory phase of mating, we noted that approximately 64% of female-approach bouts led to SI, whereas 24% led to mounting (**Figure 2C**), together accounting for the majority of SI and mount events. At the single-unit level, while some units were activated across all three behaviors (approach, SI, and mount), there were also distinct subpopulations that were selectively activated during either SI or mounting (Figure S2H). Specifically, among the 24 units activated during mounting, 11 were also co-activated during SI (**Figure 2D**), though their firing rates were significantly higher during mounting than during SI (Figure S2I). The remaining 13 neurons were selectively activated during mounting. These observations suggest that comparing the activity of single units around SI and mounting may provide a natural contrast for dissecting mount-specific mPOA neural dynamics that drive the transition to consummatory mating.

To further explore this, we aligned the activity of mount-activated units to the onset of SI and mounting. A clear inhibitory phase emerged in a window preceding mounting but was absent before SI (**Figure 2E**). To determine whether this pre-mount inhibition predicted subsequent activation during mounting, we compared each unit’s trough activity during the inhibitory window to its peak activity during mounting. This analysis revealed a significant negative correlation (**Figure 2F**, p < 0.01): the stronger the pre-onset inhibition, the greater the subsequent activation. Notably, this pre-onset suppression was prominent in neurons selectively activated during mounting (**Figure 2G**) but absent in neurons specifically activated during SI (Figure S2J), suggesting that inhibitory inputs may selectively gate the recruitment of mPOA neurons for mounting.

*Ex vivo* studies have shown that a subset of mPOA neurons exhibit post-inhibitory rebound firing (PIR)¹² following hyperpolarization. The timing of *in vivo* pre-mount suppression aligns with the conditions that induce PIR (**Figure 2E** **& 2G**), suggesting PIR may drive mPOA activation during mounting. Consistently, many mount-activated units exhibited a characteristic PIR firing pattern — prolonged suppression followed by a rapid increase in firing (**Figure 2H**). Thus, PIR, rather than disinhibition, may represent the circuit mechanism by which inhibitory input primes mPOA neurons for consummatory mating.

### Biophysical modeling PIR in mPOA neurons

To understand how synaptic inhibition could promote mPOA activation via PIR, we developed a biologically plausible single-neuron mathematical model^16–20^ (**Figure 3A**). In this model, the membrane potential V(t) of mPOA neuron was governed by synaptic inputs – modeled as the difference between excitatory and inhibitory currents^5,7^(𝐼_!"#_(𝑡) = 𝐼_!"#_(𝑡) − 𝐼_$%&_(𝑡)) – and intrinsic ionic currents mediated by Na^+^, K^+^, HCN, and T-type Ca^2+^ channels^20^. Each ionic current was controlled by voltage-dependent gating variables (e.g. *m, h, n, p, q, o*), whose dynamics follow first-order kinetics: 𝑑𝑥(𝑡)⁄𝑑𝑡 = (𝑥_’_(𝑉) − 𝑥(𝑡))⁄𝜏_"_(𝑉), where 𝑥_’_(𝑉) and 𝜏_"_(𝑉) denote the steady-state value and time constant, respectively. The values of activation and inactivation gating variables for Na^+^ (*m, h*) and T-type Ca^2+^ channels (*p, q*) are plotted in **Figure 3B-3C**, with the full equations for all ion channels shown in Table S1. Each ionic current was further defined by a reversal potential 𝐸_"_ and a maximum conductance value 𝑔_"_, chosen within physiologically realistic range (Table S2). As such, the T-type Ca^2+^ current was modeled as 𝐼_(_(𝑡) = 𝑔_(_𝑝^)^𝑞(𝑉(𝑡) − 𝐸_(_), where *p* and *q* represent activation and inactivation gating variables^20,21^, and *ET* is the Ca^2+^ reversal potential.

**Figure 3.**
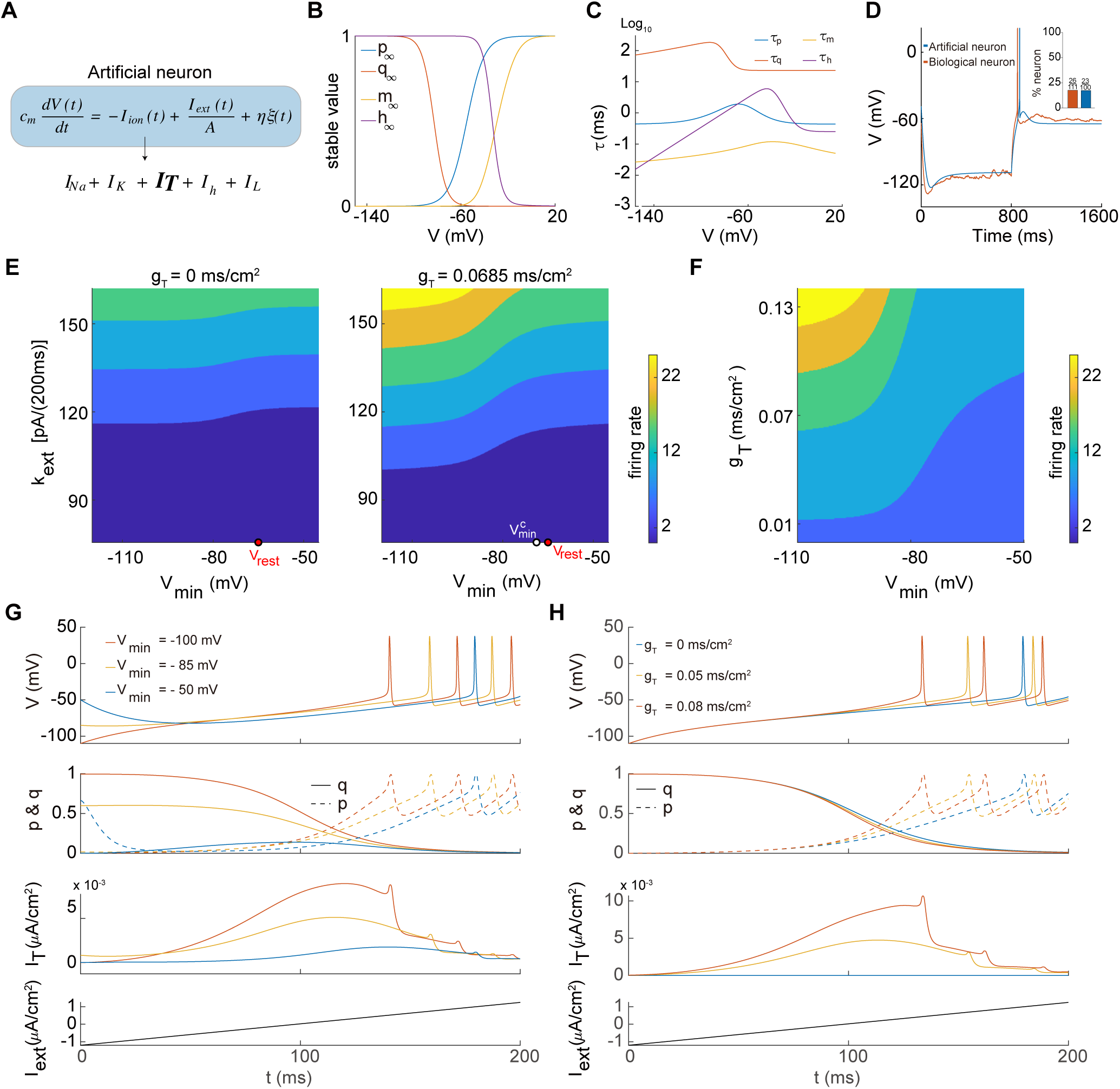
Synaptic inhibition and T-type Ca²⁺ currents interact to drive PIR in mPOA neurons. (A) Artificial mPOA neuron model incorporating sodium (*INa*), delayed-rectifier potassium (*IK*), T-type calcium (*IT*), HCN (*Ih*), and leak (*IL*) currents. (B) Stable-state activation/inactivation variables for T-type Ca^2+^ channels (𝑝_’_, 𝑞_’_) and Na^+^ channels (𝑚_’_, ℎ_’_) across different membrane potentials. (C) Time constants of activation/inactivation variables for T-type Ca^2+^ (𝜏_5_, 𝜏_4_) and Na^+^ channels (𝜏_*_, 𝜏_&_) across membrane potentials. (D) Rebound firing in biological (red) and artificial (blue) mPOA neurons in response to a hyperpolarizing current pulse (biological neuron, -100 pA; artifical neuron, -99 pA). Inset: comparable proportion of biological vs. artificial neurons exhibiting rebound firing. (E) Phase diagrams of firing rates with 𝑔_!_ = 0 mS/cm² (left) or 𝑔_!_ = 0.0685 mS/cm² (right), under varying hyperpolarization ( 𝑉_!"#_) on the X axis and excitatory input ramp (𝑘_$%&_) on the Y axis. The white dot marks the minimum 𝑉_!"#_ required to produce PIR; the red dot marks the resting membrane potential ( 𝑉_’$(&_). Color scale: firing rate (Hz). (F) Phase diagram of firing rates under varying 𝑉_!"#_and 𝑔_!_with a fixed 𝑘_$%&_ = 132 𝑝𝐴/200𝑚𝑠. **(G-H)** Simulation membrane potential traces (top) under the indicated excitatory input (bottom) with the corresponding T-type Ca^2+^ channel activation (*p*) and inactivation (*q*) variables (2^nd^ row) and *IT* amplitude (3^rd^ row). (G) Effect of different initial hyperpolarization depth (color-coded) when 𝑔_!_= 0.0685 mS/cm². (H) Effect of varying 𝑔_!_ (color-coded) under fixed hyperpolarization 𝑉_!"#_ = −110𝑚𝑉. *See also Figures S3, Table S1-S3*.

We first validated the model by simulating firing response of the neuron to depolarizing current steps using a fixed parameter set (Table S2). The resulting voltage traces closely matched traces of previous experimental recordings^12^ (Figure S3A-S3B). To capture individual variability observed in neurons, we generated 100 random parameter sets within physiologically defined ranges and simulated neuronal responses. We then assessed the mean and variability of 13 electrophysiological features, including resting membrane potential, rheobase, action potential shape, and PIR-related metrics (See Supplementary Note 1 for details). The simulated outputs showed strong agreement with experimental values (Table S3).

Critically, applying inhibitory inputs (−99 pA, 800 ms) to the modeled neuron induced a rapid hyperpolarization of the membrane potential that stabilized at a minimum value (𝑉_*$%_), consistently below the resting potential (𝑉_+!,#_) (Figure S3C). The magnitude of 𝑉_*$%_ scaled with the strength of the inhibitory input (Figure S3D). Upon release from inhibition, the model neuron exhibited rebound depolarization (Figure S3C) and, in a subset of simulations, generated PIR spikes (**Figure 3D**). Notably, 23 out of the 100 simulations produced PIR, closely matching the ∼ 23.4 % incidence rate observed in *ex vivo* recordings (26 out of 111 mPOA neurons). Thus, this validated model provides a robust platform to explore how synaptic inhibition contributes to activation of mPOA neurons.

### Inhibitory inputs interact with T-type Ca²⁺ currents to elevate mPOA firing via PIR

Given that mount-active mPOA neurons exhibit pre-onset inhibition followed by robust activation, we modeled their activity under two classes of input: (1) prolonged inhibitory inputs lasting several seconds, which stabilized the membrane potential at a hyperpolarized state (*Vmin*) at time t = 0, and (2) ramping excitatory drive, simplified as 𝐼_!"#_(𝑡) = 𝐼_-_ + 𝑘_./0_𝑡 , where 𝑘_./0_ represents the slope (strength) of excitatory input. These simulations were performed using a fixed set of electrophysiological parameters for the model neuron (Table S3). Here, 𝐼_-_ = −65 𝑝𝐴, and the firing rate is calculated as the frequency of APs generated by the mPOA neuron within 200ms. Based on the computational simulation design, we uniformly selected 751 different values of 𝑉_*$%_ within the range of [−120, −45] mV and 801 different values of 𝑘_./0_ within the range of [75.6, 162] pA/(200ms). For each parameter combination (𝑉_*$%_, 𝑘_./0_), we evolved the dynamical equations of mPOA neurons the calculated the corresponding firing rate.

In the absence of T-type Ca²⁺ conductance (𝑔_(_= 0 mS/cm^2^), neurons fired via Na⁺ currents. when excitatory input was strong enough to overcome preceding inhibition (**Figure 3E**, left). However, introducing a physiologically realistic T-type Ca²⁺ conductance ( 𝑔_(_ = 0.0685 mS/cm^2^) dramatically altered neuronal excitability (**Figure 3E**, right). Fundamentally, this transformation arises from the voltage-dependent gating properties and the distinct activation-inactivation kinetics of the T-type Ca²⁺ channels (**Figure 3B-3C**). During the inhibitory phase, the inactivation gate variable *q* approaches its steady-state value 𝑞_’_(𝑉_*$%_) , effectively “priming” the channel for activation. Upon sufficient depolarization that release the neuron from inhibition, the fast activation variable *p* rapidly increases, while *q* remains permissively maintained at 𝑞_’_(𝑉_123_) due to its slower kinetics (𝜏_4_ ≫ 𝜏_5_). This temporal mismatch enables a transient but substantial inward Ca²⁺ current to emerge: 𝐼_(_(𝑡) = 𝑔_(_𝑝^)^𝑞(𝑉(𝑡) − 𝐸_(_) ≈ 𝑔_(_(𝑉(𝑡) − 𝐸_(_) (**Figure 3G, 3H,** middle bottom), which amplifies depolarization and enhances neuronal excitability. As a result, excitatory inputs that would otherwise be subthreshold in the absence of T-type Ca²⁺ currents now reliably elicit spikes, while suprathreshold inputs produce even stronger firing responses (**Figure 3E**).

We next examined how, under the condition of PIR, varying the strength of hyperpolarization affected mPOA neuron firing and found that firing scaled with inhibitory strength when 𝑔_(_ and excitatory input were held constant above a critical threshold. Stronger inhibition, reflected by a lower 𝑉_*$%_ , resulted in earlier firing onset and higher firing rates (**Figure 3F-3G**). This aligns with our *in vivo* observation that the magnitude of the pre-onset suppression negatively correlated with the activation of mount-activated units (**Figure 2F**; **Figure 3F-3G**). To further assess how T-type conductance modulates PIR and mPOA activity, we systematically varied 𝑔_(_ while holding inhibitory and excitatory inputs constant. We found that increasing 𝑔_(_ accelerated firing onset and enhanced firing rates (**Figure 3F, 3H**). Together, these simulations demonstrate that synaptic inhibition and T-type Ca²⁺ channels act in concert to drive rebound excitation in mPOA neurons, establishing PIR as a mechanism linking inhibitory input to mount-relevant activation.

### Cacna1g is essential for PIR and male mating

Our model predicted a critical role for T-type Ca²⁺ currents in PIR–dependent mPOA activation and male mating progression. To experimentally validate this, we analyzed published single-cell transcriptome data from the hypothalamus^22^, focusing on the expression of three T-type calcium channel subunits: *Cacna1g* (*Cav3.1*), *Cacna1h* (*Cav3.2*), and *Cacna1i* (*Cav3.3*). Among these, *Cacna1g* showed the highest expression in the mPOA (Figure S4A-S4C). *In situ* hybridization confirmed robust *Cacna1g* mRNA expression in male mPOA, which was significantly reduced by castration (Figure S4D-S4E), suggesting hormonal regulation potentially relevant to male mating behaviors.

To test the functional importance of *Cacna1g*, we injected lentivirus encoding either shRNA against *Cacna1g* or a scrambled control sequence into the mPOA (**Figure 4A-4B**). The shRNA effectively reduced *Cacna1g* expression compared to controls (**Figure 4C-4D**, Figure S5A). *Ex vivo* whole-cell recordings from acute brain slices revealed that PIR was virtually abolished following *Cacna1g* knockdown (**Figure 4E- 4F**). Specifically, only 1 of 22 neurons showed rebound firing in the knockdown group, compared to 3 of 12 neurons in the scramble controls and 26 of 111 neurons previously reported in wildtype males^12^(**Figure 4F**). These data demonstrate that *Cacna1g* expression is essential for PIR in mPOA neurons.

**Figure 4.**
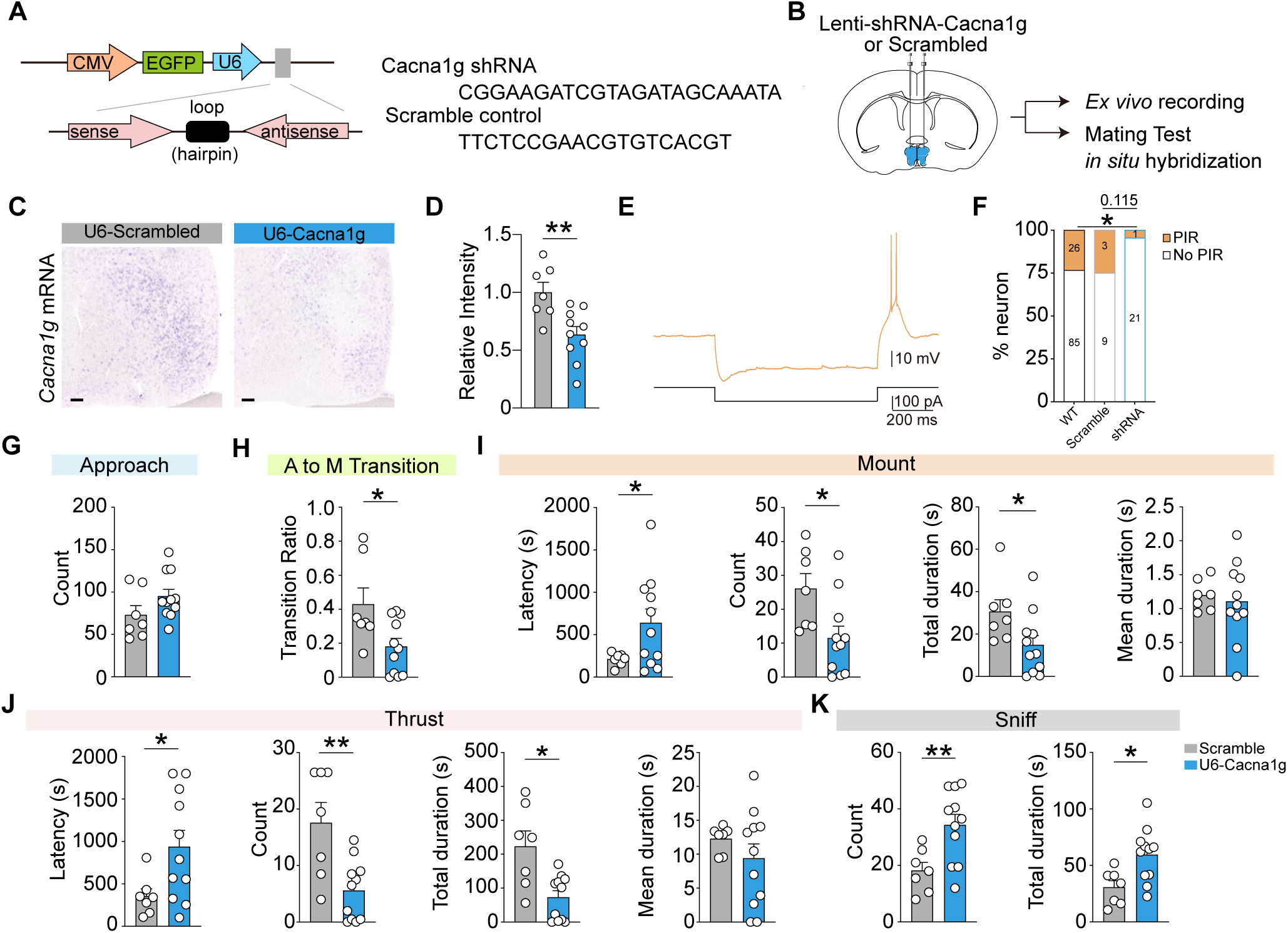
*Cacna1g* knockdown in the mPOA abolishes PIR and impairs the transition to mounting. **(A-B)** Schematics of the viral constructs (A) and experimental design (B) for lentiviral delivery of scrambled control or *Cacna1g* shRNA into the mPOA. **(C-D)** Representative *in situ* hybridization images (C) and quantification (D) showing reduced *Cacna1g* mRNA expression in the mPOA in shRNA-injected males compared to controls. Each circle represents an animal; data are presented as mean ± SEM. Scale bars, 100 μm. N = 7 scramble, 10 shRNA. **(E-F)** *Ex vivo* recordings of PIR (E) and summary of the proportion (F) of neurons exhibiting PIR across groups (F). WT data are replotted from previous results^12^. N = 111 cells from 14 WT; 12 cells from 2 scramble, and 22 cells from 5 shRNA male. **(G-K)** Quantification of sexual behaviors in control and knockdown males. Each circle represents one animal; data are presented as mean ± SEM. N = 7 scramble, 11 shRNA. *p < 0.05, **p < 0.01. *See also Figures S4&S5*.

To evaluate the behavioral consequences of impaired PIR, we bilaterally knocked down *Cacna1g* in the mPOA and assessed male mating behavior (**Figure 4B**). Knockdown males approached females as frequently as controls, indicating intact social motivation (**Figure 4G**). However, they were significantly impaired in initiating mounting and pelvic thrusting, exhibiting longer latencies and reduced frequency and total duration in these behaviors (**Figure 4H-4J**). Once initiated, however, the average duration of individual mount and pelvic thrust bouts was unchanged, suggesting normal motor execution (**Figure 4I-4J**). Interestingly, knockdown males showed increased frequency and duration of appetitive anogenital sniffing (**Figure 4K**), consistent with impaired transition from appetitive to consummatory mating. Importantly, circulating testosterone levels were unaffected by *Cacna1g* knockdown (scramble vs. shRNA: 2.42 ± 1.59 vs. 3.17 ± 1.76 ng/mL, p = 0.33). Together, these results indicate that *Cacna1g*, critical for PIR, is essential for consummatory mating.

### Cacna1g overexpression facilitates male mating behavior

Our computational model also predicted that enhancing T-type Ca²⁺ currents would promote mPOA activation and male mating. Supporting this idea, *post-hoc* analysis in wildtype males revealed a strong negative correlation between endogenous *Cacna1g* levels and mount latency (Figure S4F-S4G, r = - 0.89, p < 0.05), implicating *Cacna1g* in facilitating mating. To test causality, we used a CRISPRa-based system with a VP64 activator domain^23^ to selectively upregulate *Cacna1g* expression in the mPOA (**Figure 5A**). *In situ* hybridization confirmed robust *Cacna1g* overexpression relative to controls (**Figure 5B-5D**, Figure S5B). Behaviorally, *Cacna1g* overexpression did not change the total number of mounts in males but significantly reduced mount latency and improved the efficiency of transition from approach to mounting (**Figure 5E-5H**). Sniff frequency and duration also decreased (**Figure 5I**), indicating an accelerated shift from appetitive to consummatory mating in *Cacna1g* overexpression males. As a result, ejaculation latency was also significantly shortened (**Figure 5J**). These findings support the model that *Cacna1g*-dependent PIR facilitates mPOA activation and drives mating progression in males.

**Figure 5.**
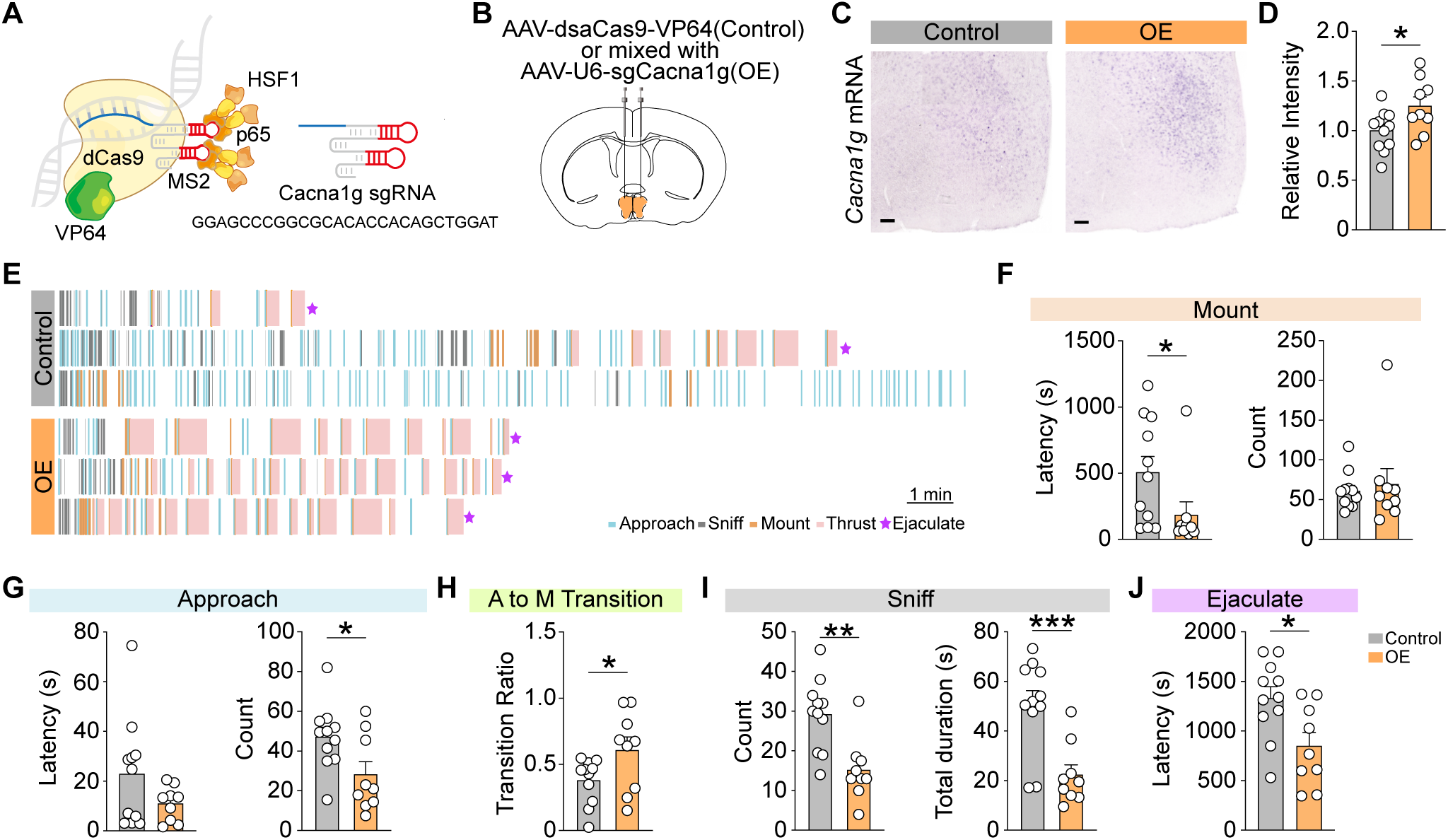
*Cacna1g* overexpression in the mPOA facilitates male mating. **(A-B)** Schematics of the VP64-based system (A) and experimental design (B) for mPOA injection with AAV-dsaCas9-VP64 alone (control) or in combination with AAV- U6-sgCacna1g (overexpression, OE). **(C-D)** Representative *in situ* hybridization images (C) and quantification (D) showing increased *Cacna1g* mRNA expression in the mPOA of OE males compared to controls. Scale bars, 100 μm. Each circle represents one animal; data are presented as mean ± SEM. N = 11 control, 9 OE. **(E)** Example raster plots from representative control and OE males during mating tests, showing behavioral events including approach (cyan), sniff (gray), mount (orange), thrust (peach), and ejaculation (magenta asterisks). OE males exhibit accelerated mating. Each row depicts the mating trial of a single male; scale bar, 1 min. **(F-J).** Quantification of sexual behaviors in control and OE males. Each circle represents one animal; data are presented as mean ± SEM. N = 11 control, 9 OE. *p < 0.05, **p < 0.01, ***p < 0.001. *See also Figures S4&S5*.

## Discussion

Unlike cortical circuits, subcortical limbic regions such as the hypothalamus are dominated by inhibitory connections^24–27^. Inhibitory inputs from regions such as the BNSTpr convey sensory and motivational signals to hypothalamic nuclei to control innate behaviors including mating and aggression in a sex-specific manner^5,6,8,10,27–30^. A long-standing hypothesis posits that these inhibitory inputs drive hypothalamic activation through disinhibition^6,7^ — a mechanism well-established in other circuits, such as the disinhibitory control of dopamine neurons in the ventral tegmental area (VTA)^31–33^ and the descending pain pathways in the periaqueductal gray (PAG)^34^. However, recent *ex vivo* studies have revealed sparse local connectivity in many hypothalamic nuclei^15^, raising questions about whether disinhibition is a viable mechanism for inhibition-driven activation in this context.

Here, we present data that support post-inhibitory rebound firing (PIR) as a distinct mechanism by which inhibition directly promotes activation in hypothalamic neurons. Rather than disinhibiting local interneurons, upstream input directly hyperpolarizes mPOA neurons in a temporally structured pattern that predicts their subsequent firing. Using a biophysically grounded computational model, we demonstrate that hyperpolarization interacts with low-threshold Ca^2+^ currents to drive rebound excitation in mPOA neurons. Experimentally, knockdown of *Cacna1g* nearly abolishes PIR and significantly impairs mating, while its overexpression enhances mating behavior. These results provide converging theoretical and empirical evidence that PIR is a key mechanism for inhibition-dependent hypothalamic activation.

Notably, both inhibitory inputs and T-type Ca²⁺ currents show male-biased properties in the mPOA. BNSTpr neurons are more strongly activated when males interact with females than when they encounter other males^6,10^, and T-type Ca²⁺ currents are more prominent in male than in female mPOA neurons^12^. Within the PIR framework, both inhibition and T-type Ca²⁺ currents must exceed a critical threshold to drive activation. As such, these quantitative sex differences in inhibitory inputs and T-type Ca²⁺ currents may account for the qualitative divergence in mPOA activation and the display of behavior: males readily transition to consummatory mating when presented with receptive females, whereas females, despite engaging in social interactions with receptive females, rarely display mating behaviors^3^. Thus, PIR offers a plausible cellular mechanism linking sex-biased circuit properties to sexually dimorphic behavioral outcomes.

While our findings focus on the mPOA in the context of male mating, we propose that PIR may represent a broader computational motif in hypothalamic circuits. Supporting this idea, biophysical studies in the rat caudal hypothalamus revealed that many neurons exhibit PIR that depends on low-voltage-activated Ca^2+^ currents^35^. Similarly, *ex vivo* recordings from hypothalamic organotypic slices demonstrated that low concentrations of oxytocin enhance presynaptic GABA_A-mediated inhibition in the supraoptic nucleus (SON), triggering rebound firing in ∼ 40% of neurons—an effect that also relies on low-threshold Ca²⁺ channels^36^. PIR has also been implicated in aggression-related circuits: neurons in both the ventrolateral division of the ventromedial hypothalamus (VMHvl) and the ventral premammillary nucleus (PMv) exhibit rebound firing in response to inhibitory input^37–39^, contributing to aggression regulation. These examples suggest that PIR may be a general strategy by which inhibitory input drives activation in hypothalamic networks.

Computationally, PIR enables postsynaptic neurons to respond selectively to temporally structured excitatory and inhibitory inputs. In the auditory system, In the auditory system, PIR is thought to endow neurons in the inferior colliculus (IC) with the ability to detect sound features defined by precise interval^40,41^, a property that computational models incorporating PIR can replicate^42^. By analogy, innate social behaviors such as male mating and aggression involve tightly timed transitions between appetitive and consummatory phases ^2,43^, requiring coordinated integration of sensory cues, internal states, and motor output. In these contexts, PIR may serve as a circuit-level timing mechanism—linking inhibition to timed activation in ways that disinhibition alone may not achieve — thereby facilitating appropriately behavioral transitions.

Beyond timing, PIR is known to also promote pacemaker-like and oscillatory dynamics^44,45^. The mPOA and BNSTpr —reciprocally connected inhibitory regions^28^— may leverage this property: BNSTpr inputs, conveying sensory cues and motivational drives, could trigger mPOA activation via PIR, while mutual inhibition between these two regions may give rise to rhythmic, sustained activity that organizes consummatory mating sequences. In summary, our results identify PIR as a key mechanism by which inhibition promotes hypothalamic activation to initiate male mating. More broadly, these findings shed light on how inhibitory limbic circuits may encode timing, behavioral transitions, and motivational drives, with potential analogs in artificial neural networks^46^.

## Resource availability

### Lead Contact

Further information and requests for resources and reagents should be directed to and will be fulfilled by the Lead Contact, Xiaohong Xu.

### Materials Availability

All unique/stable reagents generated in this study are available from the Lead Contact upon request.

### Data and Code Availability

This paper analyzes existing, publicly available data, accessible at https://doi.org/10.17863/CAM.87955
All data reported in this paper will be shared by the lead contact upon request.
This paper does not report original code.
Any additional information required to reanalyze the data reported in this paper is available from the lead contact upon request.

## Acknowledgments

We thank members of the Xu Lab for comments on the manuscript, and thank members of NPC, and Drs. Gu Yu and Jianguang Ni for valuable discussions. This work was supported by the National Science and Technology Innovation 2030 Major Program (2021ZD0203200-03 to X.X.); the National Science Foundation of China (31900721 to W.Z.); the Shanghai Municipal Science and Technology Major Project (2018SHZDZX05 to X.X.); and the Lingang Laboratory (LG202104-01-04 to X.X.).

## Author contributions

Conceptualization: W.Z. X.Z. and X.X.; Paper writing: W.Z., X.Z., L.-P.G, Y.-Y.M. and X.X.; Experiments: W.Z., L.F., X.-Z.C., Z.-L.J, S.-S.L., Z.-Y.W., and H.-T.X.; Experimental data analysis: W.Z., X.Z., Z.-W.Z., Y.-X.B., T.-M.Y., X.-J.D. and X..X.; Model construction and analysis: L.-P.G. Z.-T.F. and Y.-Y.M.

## Declaration of generative AI and AI-assisted technologies

During the preparation of this work, Xiaohong Xu used ChatGTP/OpenAI in order to improve the readability of the text. After using this tool/service, the authors reviewed and edited the content as needed and take full responsibility for the content of the publication.

## STAR Methods

### Key Resources Table

| REAGENT or RESOURCE | SOURCE | IDENTIFIER |
| --- | --- | --- |
| Bacterial and virus strains |  |  |
| Lenti-CMV-eGFP-U6-<br>*Cacna1g*-shRNA | This paper | N/A |
| Lenti-CMV-eGFP-U6-Scramble-shRNA | This paper | N/A |
| AAV-*Cacna1g*-sgRNA-MS2/P65/HSF1-U6 | This paper | N/A |
| AAV-CMV-dSaCas9/VP64 | This paper | N/A |
| Chemicals, peptides, and recombinant proteins |  |  |
| 17β-estradiol benzoate (E2) | Sigma-Aldrich | Cat #E8875 |
| Progesterone | Sigma-Aldrich | Cat #P0130 |
| DAPI | Sigma-Aldrich | Cat #D9542 |
| Experimental models: Organisms/strains |  |  |
| Mouse: C57BL/6 | Slac Laboratory Animal | N/A |
| Critical commercial assays |  |  |
| Testosterone ELISA kit | DRG Instruments GmbH | Cat#EIA-1559 |
| Software and algorithms |  |  |
| GraphPad Prism 10 | GraphPad | https://www.graphpad.com/scientific-software/prism/; RRID: SCR_002798 |
| Offline sorter V3 | Plexon | https://plexon.com/products/offline-sorter |
| pCLAMP 11 | Molecular Devices | https://www.moleculardevices.com/; RRID: SCR_011323 |
| Fiji (ImageJ) | NIH | https://imagej.nih.gov/ij/index.html; RRID: SCR_003070 |
| Adobe Illustrator CC | Adobe | https://www.adobe.com/; RRID: SCR_010279 |
| MATLAB | MathWorks | https://www.mathworks.com/products/matlab.html;<br>RRID: SCR_001622 |

### EXPERIMENTAL MODEL AND STUDY PARTICIPANT DETAILS

#### Animals

Eight-week-old adult C57BL/6 male and female mice were obtained from the Slac Laboratory Animal (Shanghai). Male and female were separated housed in the Institute of Neuroscience animal facility on 12h light/dark cycle and with food and water *ad libitum* until use. All experimental protocols were approved by the Animal Care and Use Committee (IACUC No. NA-016-2016).

### METHOD DETAILS

#### Virus

*Cacna1g* targeting sequence CGGAAGATCGTAGATAGCAAATA and control scramble sequence TTCTCCGAACGTGTCACGT, were cloned into vector pLKD- CMV-eGFP-U6-shRNA and packaged in lentivirus by Obio Technology Co. (Shanghai). Lentivirus was diluted to 1×10^8^ TU/mL in titer before injection.

The CRISPR-Cas9 over-expression system was designed and tested *in vitro* by Suzhou Haixing Biosciences Co., Ltd. The *Cacna1g* targeting sgRNA sequence were GGAGCCCGGCGCACACCACAGCTGGAT. The adeno-associated virus (AAV) containing sgRNA-MS2/P65/HSF1-U6 (2.4×10^13^ GC/mL), and another AAV containing CMV-dSaCas9/VP64 (9.14×10^13^ GC/mL) were combined at 1:1 ratio before injection to selectively upregulate *Cacna1g* expression.

#### Stereotactic Surgery

Adult mice were anesthetized with isoflurane (0.8-5 %) and placed onto a stereotactic apparatus (David Kopf Instrument, Model 1900). The skull was exposed with a small incision and holes were drilled for virus delivery with glass pipettes (15-25 μm in diameter at the tip). The coordinates of mPOA were, bregma: AP, +0.000 mm; ML, ±0.450 mm; DV, -5.100 mm (Paxios and Franklin mouse brain atlas, 2nd edition).

For bilateral virus injection, 200–400 nL of virus was injected per site at a flow rate of 100 nL/min using a homemade nanoliter injector. Glass pipettes were left in place for about 10 min post-injection before being slowly withdrawn. Mice were allowed to recover and express the virus for at least 3 weeks prior to behavioral testing.

For electrode implantation, a custom-made microdrive array consisting of 16 tungsten electrodes (diameter: 23 μm; travel length: 4 mm; Innovative Neurophysiology, Inc.) was unilaterally implanted into the mPOA at a DV coordinate of -4.700 mm. A Silver ground wire was affixed to the skull screw over the cerebellum. The entire microdrive array was secured onto the skull with dental cement (Super-Bond C&B). Mice were allowed to recover for a minimum of 5 days before *in vivo* recoding and behavioral testing.

#### Female receptivity induction

Female mice were surgically ovariectomized and supplemented with hormones as described previously^3^. Briefly, hormones were suspended in sterile sunflower seed oil (Sigma-Aldrich, S5007). To induce receptivity in females, we injected 10 μg (in 50 μL oil) and 5 μg (in 50 μL oil) of 17β-estradiol benzoate (E2) (Sigma-Aldrich, E8875) 48 h and 24 h preceding the test respectively. On the day of the test, we injected 50 μg of progesterone (Sigma-Aldrich, P0130; in 50 μL oil) 4–6 h prior to the test.

#### Hormone assays

Trunk blood was collected at the time of sacrifice before perfusion. Serum was prepared. Hormone titers were assayed using a testosterone ELISA kit (DRG Instruments GmbH, Germany, Division of ARG international, Inc, Cat#EIA-1559) according to the manufacturer’s protocols

#### *In situ* hybridization

PCR products for generating *in situ* probes were amplified using the following primer set: *Cacna1g*, 5’- CTGCGTCAGAAGCCCAGTG -3’ and 5’- GGTGAGATGACGGTGTTGTAGC -3’. RNA probes were transcribed with T7 RNA polymerase (Promega, Cat# P207E) and digoxigenin (DIG)-labeled nucleotides. Animals were anesthetized with 1% pentobarbital sodium (50 mg/kg, i.p.; Sigma, Cat#P3761) and perfused with DEPC-treated PBS (D-PBS) followed by ice-cold 4% paraformaldehyde (PFA) in D-PBS. Afterward, brains were sectioned at 40 μm thickness using a vibratome (VT1000S; Leica). Brain sections were washed in 2×SSC buffer containing 0.1% triton for 30 min, acetylated in 0.1 M triethanolamine (pH 8.0) with 0.25% acetic anhydride (vol/vol) for 10 min, equilibrated in pre-hybridization solution for 2 h at 65 ℃ and subsequently incubated with 0.5 μg/ml of specific RNA probes in hybridization buffer overnight at 65 ℃. The next day, sections were rinsed in prehybridization solution and pre-hyb/TBST (TBS with 0.1% tween-20) for 30 min each. Next, sections were washed with TBST for twice and TAE for three times, each for 5 min. Sections were then transferred into wells made of 2% agarose gel, which were run in 1×TAE at 60 V for 2 h to remove unhybridized probes. Sections were then washed twice in TBST, and subsequently incubated with sheep anti-digoxygenin-AP (1:2000, Roche, Cat# 11093274910) in 0.5% blocking reagent (Roche, 11096176001) at 4 °C overnight. On the second day, for brightfield staining sections were washed and stained with NBT (Roche, Cat# 11383213001) and BCIP (Roche, Cat# 11383221001) for 5–7 h at 37℃. All sections were washed in PBS after staining and mounted on glass slides. Images were captured using a VS120 microscope (Olympus) with a 20× objective for quantitative intensity analysis and a 10× objective for representative images. Signal intensity is calculated as the mean intensity of mPOA region minus its respective background. Relative intensity is normalized by dividing the mean of signal intensities from one group (sham-castrated male group, U6-Scramble group or Cas9- VP64 group) in the same batch.

#### DAPI fluorescent immunostaining

Brain sections (40 μm thickness) were stained DAPI (5 mg/mL, 1:1000, Sigma) in AT (0.1% Triton and 2 mM MgCl2 in PBS) for 10 min at room temperature. After several washes in PBS, brains sections were mounted onto glass slides. Images were captured under a 10× objective using a VS120 microscopy (Olympus).

#### Behavioral assays

Prior to shRNA virus injection, adult male mice (∼2 months old) were single-housed and paired overnight with hormonally primed ovariectomized C57BL/6 female mice twice, with one-week intervals between pairings. These training pairings were conducted prior to virus injection for shRNA but not overexpression experiments. All behavioral testing began at least 4 hours after the onset of the dark cycle.

For mice injected with shRNA or overexpression virus, behavioral testing was conducted following a 5 min habituation period in the testing chamber equipped with an infrared camera. A hormonally primed ovariectomized C57BL/6 female mouse was then introduced into the home cage, and behaviors were videotaped for ∼30 min at 25 Hz. Each animal was tested once every week for 2 trials. Post-behavioral analysis, mPOA brain sections were processed for either DAPI immunostaining (1/2 of sections only for shRNA test) or *Cacna1g in situ* hybridization (remaining 1/2). Inclusion criteria required visible GFP expression confirming successful viral transduction. All mice injected with shRNA or control virus met these criteria were included in the analysis.

Videos were manually annotated frame-by-frame using a custom written MATLAB program as previously described^3^, by experimenters blinded to experimental conditions. Behaviors were scored according to the following criteria: “social investigation” was defined as nose contact made by the resident to any part of the female’s body. “Sniff” was specifically scored when nose-to-urogenital contact occurred. “Mount” was scored when the male placed both forelimbs on the female’s back and climbed on top. “Thrust” was defined as rhythmic pelvic movements following mount. “Approach” was scored when the male moved toward the female from a distance greater than one body length, terminating upon initiation of social investigation, sniff, mount, or cessation of forward movement with direction change.

#### *Ex vivo* electrophysiology

For paired recordings, we transcardially perfused adult male mice firstly and sliced the brains coronally or sagittally in 300 μm using a Microtome (Precisionary Instruments, VF300-0Z model) in the ice-cold NMDG-ACSF solution containing (in mM) 93 NMDG, 2.5 KCl, 1.25 NaH2PO4, 30 NaHCO3, 20 HEPES, 25 Glucose, 5 sodium ascorbate, 2 Thiourea, 3 sodium pyruvate, 12 N-Acetyl-L-cysteine, 10 MgSO4, and 0.5 CaCl2 (pH 7.3-7.4). Slices were recovered in the NMDG-ACSF solution at 37 ℃ for 25 minutes, gradually adding appropriate Na^+^ spike-in solution (2 M NaCl in NMDG- ACSF in 5-min intervals) and then transferred to HEPES-ACSF containing (in mM) 92 NaCl, 2.5 KCl, 1.2 NaH2PO4, 30 NaHCO3, 20 HEPES, 25 Glucose, 5 sodium ascorbate, 2 Thiourea, 3 sodium pyruvate, 2 MgSO4.7H2O, 2 CaCl2. 2H2O at room temperature for another 1 hour of recovery before recording^47^

During recordings, slices were immersed with oxygenated artificial cerebrospinal fluid (ACSF) containing (in mM) 126 NaCl, 3 KCl, 26 NaHCO3, 1.2 NaH2PO4, 10 D-glucose, 2.4 CaCl2.2H2O, and 1.3 MgCl2.7H2O (300–310 mOsm) at 30 ℃. We used an upright fluorescent microscope (Olympus, BX51WI) with infrared DIC and a CCD camera to visualize cells. Glass pipettes (4-15 MΩ) with the internal solution containing (in mM) 126 K-gluconate, 2 KCl, 2 MgCl2.7H2O, 10 HEPES, 0.2 EGTA, 4 Na2ATP, 0.4 Na3GTP, and 10 K-phosphocreatine and 0.5% neurobiotin (Vectorlabs, #SP-1120-50) were used to patch cells. Recordings were performed using Axon Multiclamp 700B amplifiers, Digidata 1550B, and pCLAMP 10.2 (Molecular Devices).

In paired recordings, up to four neurons were simultaneously recorded. Target cells were randomly selected from nearby population unless otherwise specified. Within each recorded cohort, neurons were sequentially depolarized via current injection (one cell per trial), while postsynaptic responses were monitored in the remaining cells. Postsynaptic potentials were analyzed using Clampfit software. Chemical synaptic connectivity was determined using two criteria: 1) at least 5 postsynaptic responses detected across 10 stimulation trials with consistent latency, and 2) response latencies ≤ 6 ms. Series resistance was maintained below 40 MΩ for all recorded neurons throughout the experiment.

For rebound recordings, adult mice were anesthetized with isoflurane, perfused transcardially with ice-cold oxygenated (95% O2/5% CO2) high-sucrose solution (in mM: 2.5 KCl, 1.25 NaH2PO4, 2 Na2HPO4, 2 MgSO4.7H2O, 213 Sucrose, 26 NaHCO3).

Coronal slices (250 μm) were sectioned in ice-cold oxygenated high-sucrose solution using a vibratome (Leica, VT1200S), and then incubated in oxygenated ACSF (in mM: 126 NaCl, 2.5 KCl, 1.25 NaH2PO4, 1.25 Na2HPO4, 2 MgSO4.7H2O, 10 Glucose, 21.5 NaHCO3, 2 CaCl2.2H2O) at 34°C for 1h.

Neurons expressing shRNA were identified by their co-expressed green fluorescence under the fluorescent microscope (Olympus, BX51WI). Patch pipettes were ∼2 μm in diameter at the tip and had impedances of 3-5 MΩ when filled with intracellular solution (in mM: 135 K-gluconate, 4 KCl, 10 HEPES, 10 sodium phosphocreatine, 4 Mg-ATP, 0.3 Na3-GTP and 0.5 biocytin; pH:7.2; 265 mOsm). Recordings were acquired using Multiclamp 700B amplifier and Digidata 1440A interface (Molecular Devices). To assess the rebound firing properties, neurons were held at -60 mV under current clamp and injected with negative current (800 ms duration) ranging from 0 to -100 pA in 10 pA increments with 20 s inter-stimulus intervals. Post-inhibitory rebound firing was quantified as the percentage of trials exhibiting action potentials within 200 ms following cessation of the hyperpolarizing current pulse (-100 pA). Neurons with access resistance 25 MΩ after formation of the whole-cell were excluded. All experiments were conducted at a temperature of ∼33 °C using a temperature controller (Warner Instrument, TC324B).

#### *In vivo* electrophysiology

On recording days, a lightweight headstage was attached to the implanted electrode assembly, and mice were allowed to move freely in their home cages. Spontaneous spiking activity (digitized at 30 kHz, band-pass filtered 250-5,000 Hz) and local field potential (LFP, digitized at 2 kHz sampling rate, low-pass filtered up to 250 Hz) were recorded simultaneously through the multichannel amplifier (Blackrock system). Electrode bundles were advanced until stable spiking activities were detected. A single channel without detectable spikes served as the reference electrode. Following a 10 min recording, a hormonally primed ovariectomized C57BL/6 female was introduced into the cage for behavioral testing. Recording duration varied from 10-30 min depending on the male’s behavioral response. For mating behavior test, if ejaculation happened, female was removed approximately 5 min post-ejaculation; otherwise, the female remained in the cage for 30 min. Between recording sessions, electrodes were lowered in 50 μm increments, followed by a minimum 2-day recovery period. All procedures were conducted during the dark phase of the light cycle. The electrode positions were verified post-hoc through electrolytic lesions (30 μA, 20 s).

#### Spike sorting and frequency analysis

All waveforms recorded from each microwire were imported in Offline Sorter V3 (Plexon Inc.) for single-unit isolation. Units were manually identified using threshold- based detection followed by principal component analysis (PCA) for cluster separation. Quality control criteria included: (1) <0.1% of spikes with inter-spike intervals (ISIs) < 1.8 ms, indicating minimal refractory period violations, and (2) signal-to-noise ratio >2. Cross correlograms between simultaneously recorded channels were examined to prevent duplicate classification of single units across overlapping electrodes.

The firing frequency (F) was calculated using 0.25-second bins. Normalized frequency was defined as (F - Fbaseline) / Fbaseline, where the Fbaseline was computed as the mean frequency during the consummatory phase, excluding periods of approach, social investigation, sniffing, mounting, and thursting.

#### Cross correlogram analysis

Statistical significance of functional connectivity was assessed using jitter-based surrogate analysis^48^. Spike timestamps were randomly perturbed within a uniform ± 5 ms window to generate 1,000 surrogate datasets while preserving overall firing patterns. Cross-correlograms were computed for each surrogate dataset over a ± 40 ms window (1ms bins). Confidence intervals were established at the 99% level (p = 0.01) using pointwise statistics across all surrogate correlograms. Global significance bands were defined by the extremal values (maximum and minimum) at each time bin across all jitter datasets. Putative monosynaptic connections were identified when cross- correlogram counts exceeded these global confidence bands within the 1-4 ms latency window, with peaks indicating excitatory connections and troughs indicating inhibitory connections.

#### Single-cell dataset analysis

To examine the expression patterns of T-type calcium channel subunit genes (*Cacna1g*, *Cacna1h*, and *Cacna1i*) in the medial preoptic area (mPOA), we reanalyzed publicly available single-cell RNA sequencing data from Lukas et al., 2022. The dataset was processed using Seurat (v4.3.0) in R. Cells were first filtered by anatomical region and sex annotations provided in the metadata. Specifically, we selected cells with the ‘Region_summarized’ annotation labeled as “medial preoptic area” and ‘Inferred_sex’ labeled as “M” (male). Subsequent analysis was performed exclusively on these male mPOA neurons. No reclustering was performed. Instead, we used the original dimensionality reduction coordinates provided by the authors (umap_scvi) to generate the UMAP visualization. Cell type annotations were adopted directly from the C25_named metadata field, corresponding to the transcriptional subtypes defined in the original study.

To enable direct comparison of expression levels across the three T-type calcium channel subunit genes (*Cacna1g*, *Cacna1h*, and *Cacna1i*), we rescaled their normalized expression values using a unified scaling approach across all cells. This standardized expression matrix was used to generate feature and dot plots, where dot size indicates the percentage of cells expressing the gene (expression > 0.1), and color intensity reflects the scaled average expression.

### Data and Statistical analysis

Data analysis was conducted by MATLAB. Data were statistical analyzed and plotted with GraphPad Prism 10. Values are presented as mean ± SEM. In all cases, N refers to the number of neurons, neuronal pairs or animals. For comparison of two groups, if the data were fitted into normal distribution (Shapiro-Wilk test) and with equal variances (F test), p values were calculated with Student’s t test (two-tailed, paired, or unpaired), whereas we used t test with Welch’s correction for data with normal distribution but unequal variances; otherwise, p values were calculated with Mann-Whitney test for unpaired data, and Wilcoxon matched-pairs signed-rank test for paired data. For correlation test, Pearson r test was used for normal distribution data, and Spearman correlation test was used for abnormal distribution data Categorical data were compared with Fisher’s exact test. Table S4 lists the statistical values for all comparisons in the figures. \**p* < 0.05, \*\**p* < 0.01, \*\*\**p* < 0.001.

## Supplementary Figure Legend

Figure S1. Local connectivity in the barrel cortex revealed by *ex vivo* recordings.

Related to Figure1.

**(A)** Schematic of paired whole-cell recordings.

**(B)** Coronal brain section highlighting the barrel cortex and mPOA.

**(C)** Representative traces from paired *ex vivo* recordings, showing synaptic connectivity between barrel cortex neurons. Blue traces indicate the averaged membrane potential in the stimulated cell (#2) with evoked action potentials; green traces indicate the averaged post-synaptic responses in the unstimulated cell (#1), held at -40 mV. The original raw traces are shown in gray.

**(D)** Quantification of monosynaptic connectivity in the 20 tested neuron pairs in the barrel cortex.

Figure S2. Single-unit recordings in the mPOA. *Related to* Figures 1 *& 2*.

**(A-B)** Example image (A) and schematics (B) showing electrode tip locations in individual animals.

**(C-G)** Averaged (left) and single unit heatmap (right) of normalized firing frequency (0.25 s bins) for units activated or inhibited during approach (C), SI (D), sniff (E), mount (F), and pelvic thrust (G). Each row represents one unit. Color scale: firing rate (Hz).

**(H)** Hierarchical clustering of single units by behavioral tuning.

**(I-J)** Normalized firing rates aligned to bout onset (time 0) for mount/SI co-activated units (N=11, I) and SI-specific units (N=17, J). Traces show responses around mount (orange) and SI (green) bouts; shaded areas, SEM. Bar plots on the right quantify firing rate changes within the gray box preceding either behavior.

Figure S3. Biophysical model reproduces the responses of mPOA neurons in whole-cell recording. *Related to Figure3, Tables S1-S3*.

**(A)** Representative traces of membrane potential changes in response to depolarizing current injection. Biological neuron: red line, 40 pA; artificial neuron: blue trace, 47 pA.

**(B)** Firing rates of the biological (red) and artificial (blue) mPOA neurons across a range of depolarizing current injections (𝐼_$%&_).

**(C)** Representative traces of membrane hyperpolarization following hyperpolarizing current injection. Biological neuron: red trace, -100 pA; artificial neuron: blue trace, - 99 pA.

**(D)** Minimum membrane potential ( 𝑉_123_ ) of biological (red) and artificial mPOA neurons (blue) in response to varying hyperpolarizing currents (𝐼_ext_).

Figure S4. Gonadal regulation of *Cacna1g* expression in the male mPOA and its negative correlation with mounting latency in wildtype males. *Related to* Figures 4*&5*.

(A) UMAP plot of male mPOA neurons classified into four major clusters.

(B) Expression patterns of *Cacna1g*, *Cacna1h*, and *Cacna1i* across mPOA neuron clusters.

(C) Dot plot showing relative expression of *Cacna1g*, *Cacna1h*, *Cacna1i* across mPOA clusters. Color intensity indicates average expression; and circle size indicates the percentage of cells expressing each gene.

(D) *In situ* hybridization of *Cacna1g* mRNA in sham-castrated versus castrated males.

(E) Quantification of *Cacna1g* expression intensity in sham-castrated (blue) and castrated (gray) males. Each point represents an individual animal; bar shows mean ± SEM. N = 8 sham, 9 castrates.

(F) Representative *Cacna1g in situ* hybridization in two individual wildtype males, corresponding to the two-colored data points in (G).

(G) Negative correlation between *Cacna1g* expression intensity and mounting latency in wildtype males. Colored points correspond to the examples shown in (F).

* p < 0.05.

Figure S5. Representative images of shRNA-mediated knockdown and Cas9a- mediated overexpression of *Cacna1g*. *Related to* Figures 4*&5*.

Each column shows data from an individual male: (A) shRNA-mediated knockdown of *Cacna1g* and (B) Cas9a-mediated overexpression of *Cacna1g*.

## Supplemental information

Table S1. Steady-state values and time constants of gating variables for different ion channels in mPOA neurons. *Related to Figure 3, Figure S3*.

Table S2. Biophysical Parameters values used in simulations. *Related to Figure 3, Figure S3*.

Table S3. Comparison of simulated and experimental electrophysiological properties of mPOA neurons. *Related to Figure 3, Figure S3*.

Table S4. Detailed statistical information for each figure panel. *Related to Figure 1-5; Figure S1-5*.

**Supplementary Note 1. Artificial mPOA neuron modeling.**

