## Supplemental Figure 1-5 for "Post-inhibitory rebound firing drives hypothalamic activation for male mating"

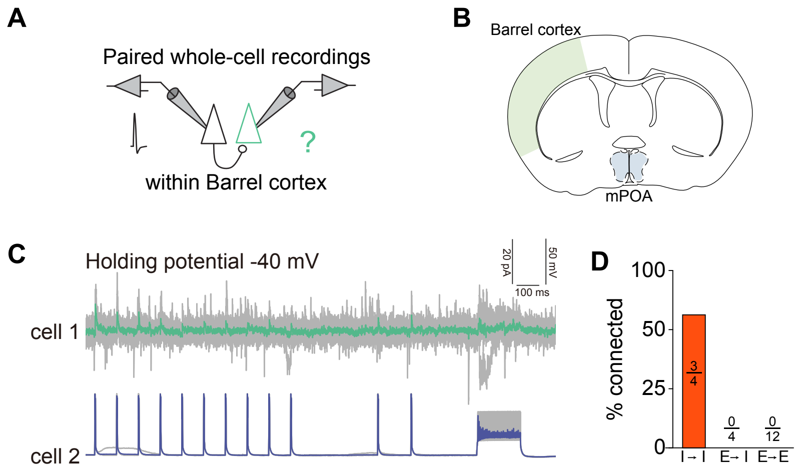


**Figure S1. Local connectivity in the barrel cortex revealed by *ex vivo* recordings. *Related to Figure1.***
**(A)** Schematic of paired whole-cell recordings.
**(B)** Coronal brain section highlighting the barrel cortex and mPOA.
**(C)** Representative traces from paired *ex vivo* recordings, showing synaptic connectivity between barrel cortex neurons. Blue traces indicate the averaged membrane potential in the stimulated cell (#2) with evoked action potentials; green traces indicate the averaged post-synaptic responses in the unstimulated cell (#1), held at -40 mV. The original raw traces are shown in gray.
**(D)** Quantification of monosynaptic connectivity in the 20 tested neuron pairs in the barrel cortex.

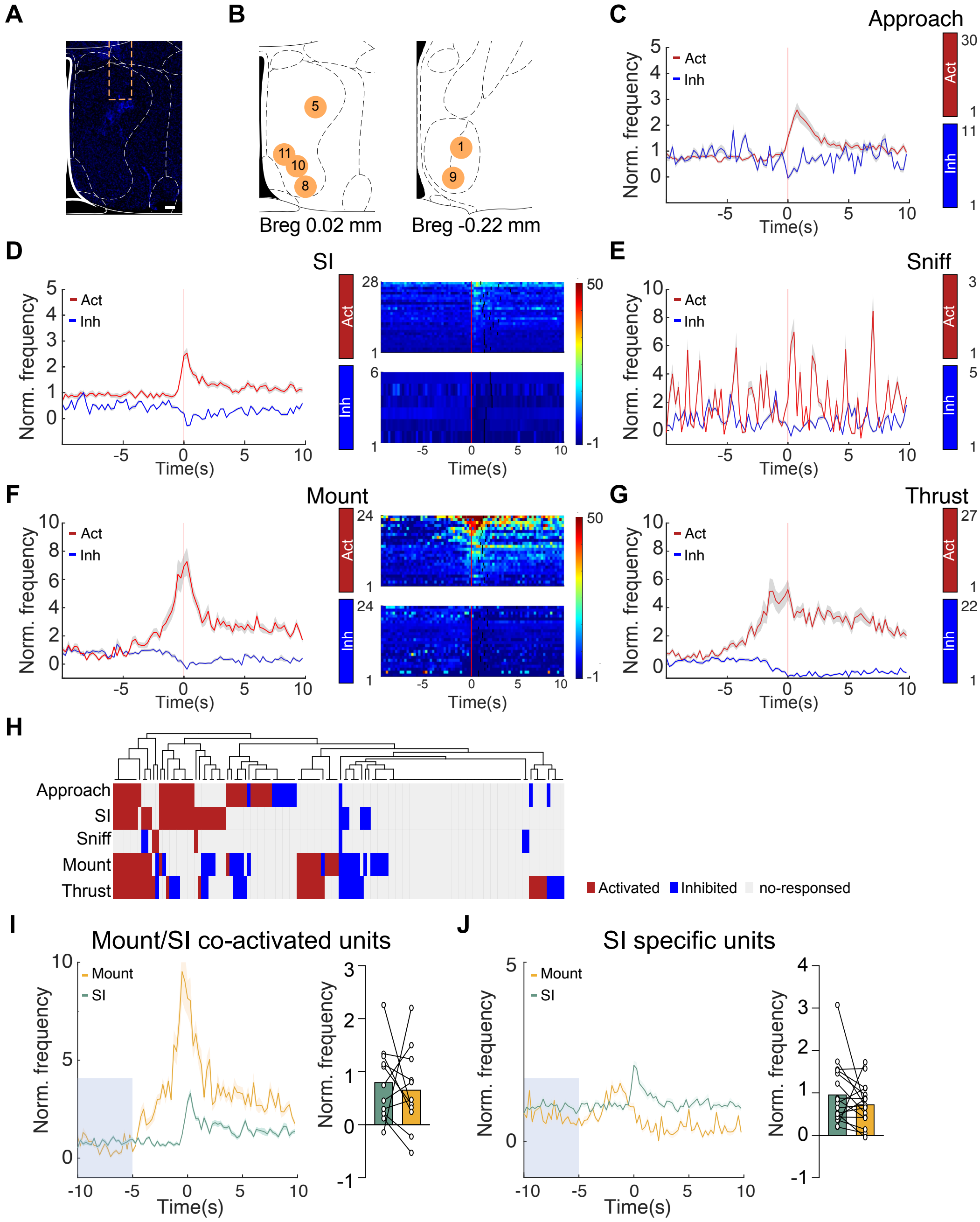


**Figure S2. Single-unit recordings in the mPOA.** ***Related to Figure1 & 2.***

**(A- B)** Example image (A) and schematics (B) showing electrode tip locations in individual animals.

**(C- G)** Averaged (left) and single unit heatmap(right) of normalized firing frequency (0.25 s bins) for units activated or inhibited during approach (C), SI (D), sniff (E), mount (F), and pelvic thrust (G). Each row represents one unit. Color scale: firing rate (Hz).

**(H)** Hierarchical clustering of single units by behavioral tuning.

**(I- J)** Normalized firing rates aligned to bout onset (time 0) for mount/SI co-activated units (N=11, I) and SI-specific units (N=17, J). Traces show responses around mount (orange) and SI (green) bouts; shaded areas, SEM. Bar plots on the right quantify firing rate changes within the gray box preceding either behavior.

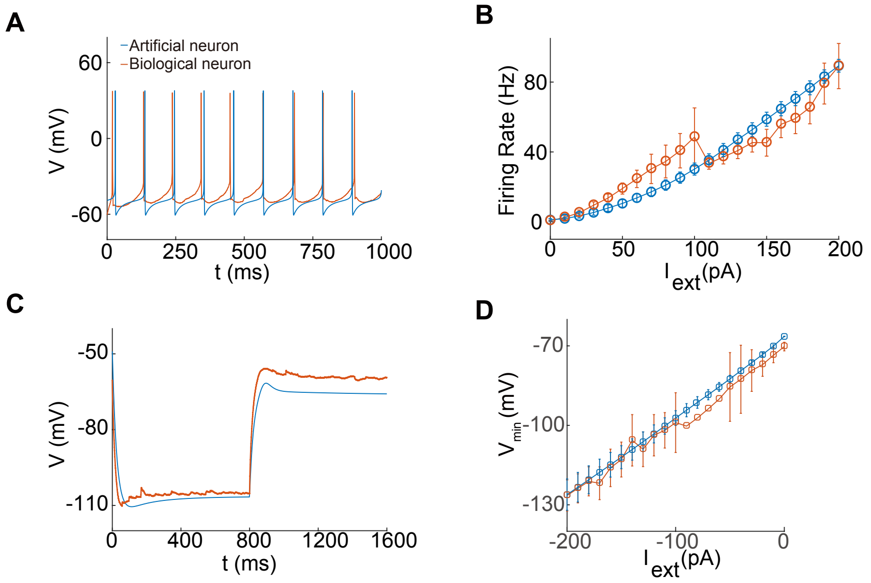


**Figure S3. Biophysical model reproduces the responses of mPOA neurons in whole-cell recording.** ***Related to Figure3, Tables S1-S3.***
**(A)** Representative traces of membrane potential changes in response to depolarizing current injection. Biological neuron: red line, 40 pA; artificial neuron: blue trace, 47 pA.
**(B)** Firing rates of the biological (red) and artificial (blue) mPOA neurons across a range of depolarizing current injections ($I_{ext}$).
**(C)** Representative traces of membrane hyperpolarization following hyperpolarizing current injection. Biological neuron: red trace, -100 pA; artificial neuron: blue trace, -99 pA.
**(D)** Minimum membrane potential ($V_{min}$) of biological (red) and artificial mPOA neurons (blue) in response to varying hyperpolarizing currents ($I_{ext}$).

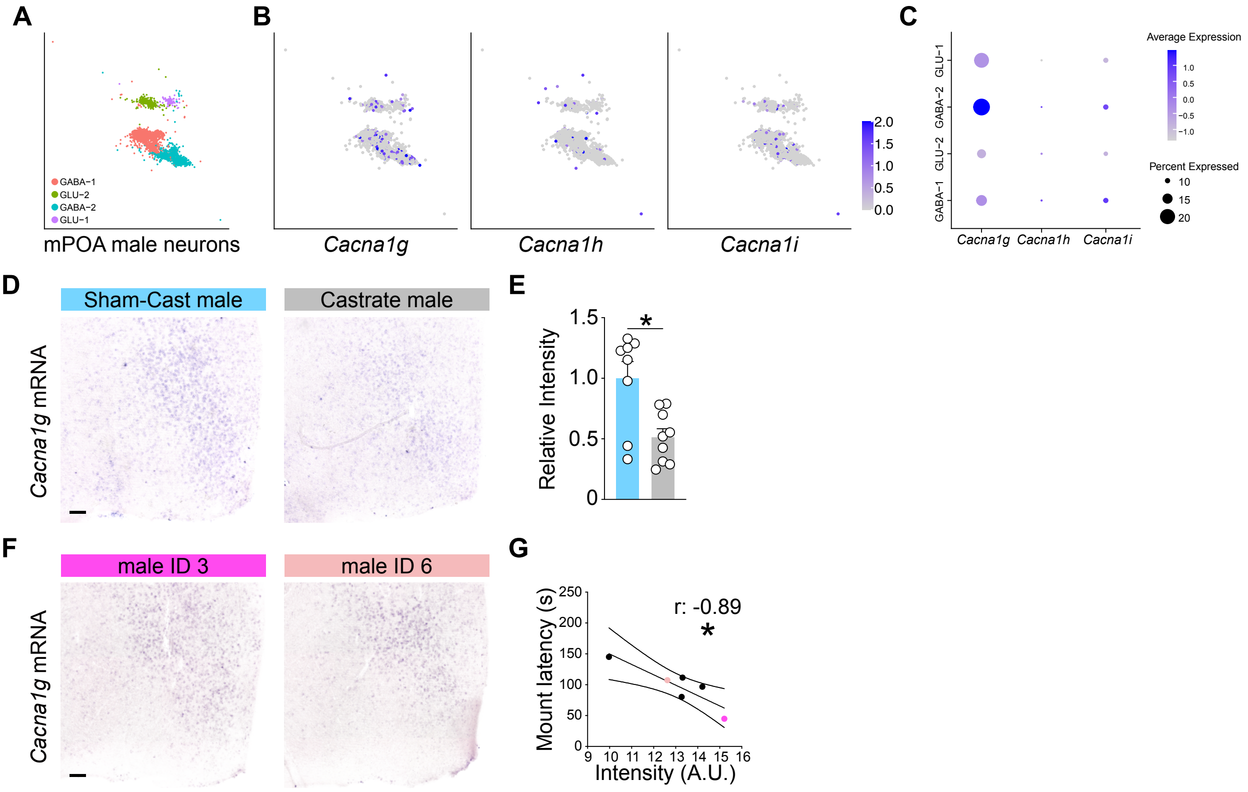


**Figure S4. Gonadal regulation of *Cacna1g* expression in the male mPOA and its negative correlation with mounting latency in wildtype males. *Related to Figures 4&5.***

**(A)** UMAP plot of male mPOA neurons classified into four major clusters.

**(B)** Expression patterns of *Cacna1g*, *Cacna1h*, and *Cacna1i* across mPOA neuron clusters.

**(C)** Dot plot showing relative expression of *Cacna1g*, *Cacna1h*, *Cacna1i* across mPOA clusters. Color intensity indicates average expression; and circle size indicates the percentage of cells expressing each gene.

**(D)** *In situ* hybridization of *Cacna1g* mRNA in sham-castrated versus castrated males.

**(E)** Quantification of *Cacna1g* expression intensity in sham-castrated (blue) and castrated (gray) males. Each point represents an individual animal; bar shows mean ± SEM. N = 8 sham, 9 castrates.

**(F)** Representative *Cacna1g* in situ hybridization in two individual wildtype males, corresponding to the two colored data points in (G).

**(G)** Negative correlation between *Cacna1g* expression intensity and mounting latency in wildtype males. Colored points correspond to the examples shown in (F).

*p < 0.05

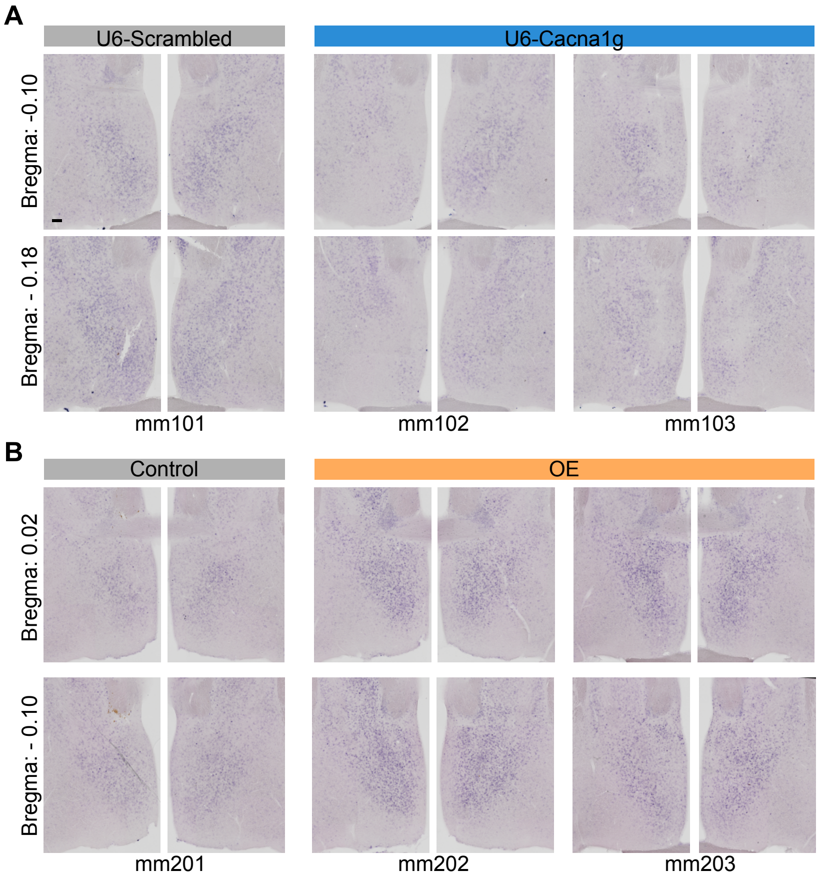


**Figure S5. Representative images of shRNA-mediated knockdown and Cas9a-mediated overexpression of *Cacna1g*. *Related to Figures 4&5.***
Each column shows data from an individual male: (A) shRNA-mediated knockdown of *Cacna1g* and (B) Cas9a-mediated overexpression of *Cacna1g*.
