## Supplementary material for "Post-inhibitory rebound firing drives hypothalamic activation for male mating": Table1

| *Variable* | *Full Name* | *Equation* |
|---|---|---|
| $\boldsymbol{m_\infty}$ | steady state of active gate of $Na^{+}$ | $m_\infty(V) = \frac{-0.32(V+42)[\exp\left(\frac{V+15}{5}\right) - 1]}{-0.32(V+42)\left[\exp\left(\frac{V+15}{5}\right) - 1\right] + 0.28(V+15[\exp\left(\frac{-V-42}{10}\right) - 1])}$ |
| $\boldsymbol{\tau_m}$ | time constant of active gate of $Na^{+}$ | $\tau_m(V) = \frac{1}{-0.32(V+42)\left[\exp\left(\frac{V+15}{5}\right) - 1\right] + 0.28(V+15[\exp\left(\frac{-V-42}{10}\right) - 1])}$ |
| $\boldsymbol{h_\infty}$ | steady state of inactive gate of $Na^{+}$ | $h_\infty(V) = \frac{0.128\exp\left(\frac{-V-38}{18}\right)}{0.128\exp\left(\frac{-V-38}{18}\right) + 4}$ |
| $\boldsymbol{\tau_h}$ | time constant of inactive gate of $Na^{+}$ | $\tau_h(V) = \frac{1}{0.128\exp\left(\frac{-V-38}{18}\right) + 4}$ |
| $\boldsymbol{n_\infty}$ | steady state of active gate of $K^{+}$ | $n_\infty(V) = \frac{-0.032(V+40)}{-0.032(V+40) + 0.5\exp\left(\frac{-V-45}{10}\right)}$ |
| $\boldsymbol{\tau_n}$ | time constant of active gate of $K^{+}$ | $\tau_n(V) = \frac{1}{-0.032(V+40) + 0.5\exp\left(\frac{-V-45}{10}\right)}$ |
| $\boldsymbol{p_\infty}$ | steady state of active gate of $Ca^{2+}$ | $p_\infty(V) = \frac{1}{1 + \exp\left(\frac{-V-55}{7.4}\right)}$ |
| $\boldsymbol{\tau_p}$ | time constant of active gate of $Ca^{2+}$ | $\tau_p(V) = \frac{3 + 1/(\exp\left(\frac{V+30}{10}\right) + \exp(\frac{-V-105}{15}))}{6.9}$ |
| $\boldsymbol{q_\infty}$ | steady state of inactive gate of $Ca^{2+}$ | $q_\infty(V) = \frac{1}{1 + \exp\left(\frac{V+83}{5}\right)}$ |
| $\boldsymbol{\tau_q}$ | time constant of inactive gate of $Ca^{2+}$ | $\tau_q(V) = \frac{85 + 1/(\exp\left(\frac{V+45}{4}\right) + \exp(\frac{-V-410}{50}))}{3.7}$ |
| $\boldsymbol{o_\infty}$ | steady state of inactive gate of *hyperpolarization - activated cation* | $o_\infty(V) = \frac{1}{1 + \exp\left(\frac{V+75}{5.5}\right)}$ |
| $\boldsymbol{\tau_o}$ | time constant of inactive gate of *hyperpolarization - activated cation* | $\tau_o(V) = \frac{1}{\exp(-14.59 - 0.086V) + \exp(-1.87 + 0.0701V)}$ |
