## Supplementary material for "Post-inhibitory rebound firing drives hypothalamic activation for male mating": Table2

| *Parameters* | *Descriptions* | *Value ($mS/cm^2$)* |
| --- | --- | --- |
| Figure 3D | | |
| $g_{Na}$ | Maximum conductance of $Na^+$ ion channel | 15.1 |
| $g_K$ | Maximum conductance of $K^+$ ion channel | 2.5 |
| $g_h$ | Maximum conductance of hyperpolarization activated cation ion channel | 0.0091 |
| $g_T$ | Maximum conductance of T-type $Ca^{2+}$ ion channel | 0.0685 |
| $g_L$ | Conductance of leaky ion channel | 0.05334 |
| Figure 3E-F | | |
| $g_{Na}$ | Maximum conductance of $Na^+$ ion channel | 15.1 |
| $g_K$ | Maximum conductance of $K^+$ ion channel | 2.5 |
| $g_h$ | Maximum conductance of hyperpolarization activated cation ion channel | 0.0091 |
| $g_L$ | Conductance of leaky ion channel | 0.038 |
| Figure 3G | | |
| $g_{Na}$ | Maximum conductance of $Na^+$ ion channel | 15.1 |
| $g_K$ | Maximum conductance of $K^+$ ion channel | 2.5 |
| $g_h$ | Maximum conductance of hyperpolarization activated cation ion channel | 0.0091 |
| $g_L$ | Conductance of leaky ion channel | 0.038 |
| $g_T$ | Maximum conductance of T-type $Ca^{2+}$ ion channel | 0.0685 |
| Figure 3H | | |
| $g_{Na}$ | Maximum conductance of $Na^+$ ion channel | 15.1 |
| $g_K$ | Maximum conductance of $K^+$ ion channel | 2.5 |
| $g_h$ | Maximum conductance of hyperpolarization activated cation ion channel | 0.0091 |
| $g_L$ | Conductance of leaky ion channel | 0.038 |
| Figure S3A | | |
| $g_{Na}$ | Maximum conductance of $Na^+$ ion channel | 15.1 |
| $g_K$ | Maximum conductance of $K^+$ ion channel | 2.5 |
| $g_h$ | Maximum conductance of hyperpolarization activated cation ion channel | 0.0091 |
| $g_T$ | Maximum conductance of T-type $Ca^{2+}$ ion channel | 0.0685 |
| $g_L$ | Conductance of leaky ion channel | 0.05334 |
| Figure S3B | | |

| | | |
|---|---|---|
| $g_{Na}$ | Maximum conductance of $Na^+$ ion channel | 15.1 |
| $g_K$ | Maximum conductance of $K^+$ ion channel | 2.5 |
| $g_h$ | Maximum conductance of hyperpolarization activated cation ion channel | 0.0091 |
| $g_T$ | Maximum conductance of T-type $Ca^{2+}$ ion channel | 0.0685 |
| $g_L$ | Conductance of leaky ion channel | 0.11 |
| Figure S3C | | |
| $g_{Na}$ | Maximum conductance of $Na^+$ ion channel | 15.1 |
| $g_K$ | Maximum conductance of $K^+$ ion channel | 2.5 |
| $g_h$ | Maximum conductance of hyperpolarization activated cation ion channel | 0.004 |
| $g_T$ | Maximum conductance of T-type $Ca^{2+}$ ion channel | 0.06 |
| $g_L$ | Conductance of leaky ion channel | 0.04 |
| Figure S3D | | |
| $g_{Na}$ | Maximum conductance of $Na^+$ ion channel | 15.1 |
| $g_K$ | Maximum conductance of $K^+$ ion channel | 2.5 |
| $g_h$ | Maximum conductance of hyperpolarization activated cation ion channel | 0.0091 |
| $g_T$ | Maximum conductance of T-type $Ca^{2+}$ ion channel | 0.0685 |
| $g_L$ | Conductance of leaky ion channel | 0.071 |
| Table S3 | | |
| $g_{Na}$ | Maximum conductance of $Na^+$ ion channel | N $(15.1,\ 5^2)$ |
| $g_K$ | Maximum conductance of $K^+$ ion channel | N $(2.5,\ 0.5^2)$ |
| $g_h$ | Maximum conductance of hyperpolarization activated cation ion channel | N $(0.0091,\ 0.0011^2)$ |
| $g_T$ | Maximum conductance of T-type $Ca^{2+}$ ion channel | N $(0.0685,\ 0.042^2)$ |
| $g_L$ | Conductance of leaky ion channel | N $(0.11,\ 0.06^2)$ |
