## Supplementary material for "Post-inhibitory rebound firing drives hypothalamic activation for male mating": Table3

| *Parameters* | *Descriptions* | *Biological cell* | *Artificial cell* |
|---|---|---|---|
| (1) $V_m$ $(mV)$ | Resting membrane potential | -70 (6.48) | -65.46 (0.62) |
| (2) $Rheobase(pA)$ | Minimum current needed to elicit a spike when holding at -60 mV | 24.18 (6.25) | 23.61(8.89) |
| (3) $Spike\ dV/dt_{max}$ $(mV/ms)$ | Maximum slope value of $V$ during the AP | 275.3 (46.7) | 249.97 (75.34) |
| (4) $V_{th}(mV)$ | The potential value when $dV/dt = 20\ V/s$ | -35.39 (2.31) | -35.11 (1.58) |
| (5) Spike peak $(mV)$ | Maximum membrane potential during the AP | 32.65 (2.31) | 33.56 (9.78) |
| (6) Spike amplitude $(mV)$ | The difference between the peak value and the threshold potential | 68.04 (5.25) | 68.67 (11.34) |
| (7) Spike half-height duration $(ms)$ | Time difference when neuron's potential reaches half of spike-peak during AP. | 0.70 (0.11) | 0.71 (0.02) |
| (8) Spike rise time $(ms)$ | Time difference between 10% and 90% of the spike amplitude during the ascending phase. | 0.31 (0.05) | 0.36 (0.05) |
| (9) Spike decay time $(ms)$ | Time difference between 10% and 90% of the spike amplitude during the descending phase. | 0.75 (0.33) | 0.53 (0.05) |
| (10) $\Delta G_{sag}(\%)$ | A sag index measuring $I_h$. | 17.93 (7.12) | 17.86 (7.12) |
| (11) $P_{rebound}$ | The probability of neurons exhibiting rebound firing after hyperpolarizing input | 23.43% | 23% |
| (12) Rebound amplitude $(mV)$ | The difference between the rebound peak and the baseline afterwards. | 3.82 (1.54) | 3.84 (1.09) |
| (13) Rebound duration $(ms)$ | The time interval between the offset of hyperpolarization to the point of rebound returning to baseline. | 232.0 (72.7) | 246.39 (44.34) |
