## Supplementary material for "Post-inhibitory rebound firing drives hypothalamic activation for male mating": Table4

### Statistical Table

| Figure | Sample Size (n) | Statistical test | p value |
|---|---|---|---|
| Figure1A-1D | Coronal sections: 53 pairs from 6 males<br>Sagittal sections: 84 pairs from 3 males | / | / |
| Figure1E-1I | 235 pairs from 6 males | / | / |
| Figure2D | Mount specific unit: 13 units<br>Mount-SI coactivated units: 11 units<br>SI specific unit: 17 units<br>Neither responded units: 87 units<br>All units were collected from 6 males | / | / |
| Figure2E | Mount activated unit: 24 units | Wilcoxon matched-pairs signed rank test | *, 0.0395 |
| Figure2F | Mount activated unit: 24 units | Spearman correlation test | R, -0.5993; **, 0.002 |
| Figure2G | Mount specific unit: 13 units | Paired t test | *, 0.0312 |
| Figure3D | 100 artificial cells<br>111 Wildtype cells | Fisher's exact test | ns, 1 |
| Figure4D | Scramble, 7 males<br>*Cacna1g*-shRNA, 10 males | Unpaired t test | **, 0.0051 |
| Figure4F | WT, 111 cells from 14 males<br>Scramble, 12 cells from 2 males<br>*Cacna1g*-shRNA, 22 cells from 5 males | Fisher's exact test | WT vs Scramble, ns, >0.9999<br>Scramble vs shRNA, ns, 0.115<br>WT vs shRNA, *, 0.0457 |
| Figure4G | Scramble, 7 males<br>*Cacna1g*-shRNA, 11 males | Unpaired t test | ns, 0.1201 |
| Figure4H | Scramble, 7 males<br>*Cacna1g*-shRNA, 11 males | Unpaired t test | *, 0.0211 |
| Figure4I | Scramble, 7 males<br>*Cacna1g*-shRNA, 11 males | Unpaired t test with Welch's correction<br>Unpaired t test<br>Unpaired t test<br>Unpaired t test with Welch's correction | Mount latency, *, 0.0267<br>Mount count, *, 0.0201<br>Mount total duration, *, 0.0441<br>Mount mean duration, ns, 0.6458 |
| Figure4J | Scramble, 7 males<br>*Cacna1g*-shRNA, 11 males | Unpaired t test with Welch's correction<br>Unpaired t test<br>Mann Whitney test<br>Mann Whitney test | Thrust latency, *, 0.0153<br>Thrust count, **, 0.0033<br>Thrust total duration, *, 0.0111<br>Thrust mean duration, ns, 0.5788 |
| Figure4K | Scramble, 7 males<br>*Cacna1g*-shRNA, 11 males | Unpaired t test<br>Unpaired t test | Sniff count, **, 0.0085<br>Sniff total duration, *, 0.0118 |
| Figure5D | Control, 11 males<br>OE, 9 males | Unpaired t test | *, 0.032 |
| Figure5F | Control, 11 males<br>OE, 9 males | Mann Whitney test<br>Unpaired t test | Mount latency, *, 0,0381<br>Mount count, ns, 0.5162 |
| Figure5G | Control, 11 males<br>OE, 9 males | Mann Whitney test<br>Unpaired t test | Approach latency, ns, 0.2299<br>Approach count, *, 0.0293 |
| Figure5H | Control, 11 males<br>OE, 9 males | Unpaired t test | Transition, *, 0.0470 |
| Figure5I | Control, 11 males | Unpaired t test | Sniff count, **, 0.0017 |

| | | | |
|---|---|---|---|
| | OE, 9 males | Unpaired t test | Sniff total duration, ***, 0.0010 |
| Figure5J | Control, 11 males<br><br>OE, 9 males | Unpaired t test | Ejaculate latency, *, 0.0150 |
| FigureS1 | Coronal sections: 12 pairs from 2 males | / | / |
| FigureS2G | Mount-SI coactivated units: 11 units | Paired t test | ns, 0.6772 |
| FigureS2H | SI specific unit: 17 units | Wilcoxon matched-pairs signed rank test | ns, 0.1594 |
| FigureS3 | 100 artificial cells | / | / |
| FigureS4E | 8 sham males<br><br>9 castration males | Mann Whitney test | *, 0.0152 |
| FigureS4G | 6 WT males | Pearson correlation test | *, 0.0170 |
